# Beyond the Genetic Code: *A Tissue Code?*

**DOI:** 10.1101/2023.03.05.531161

**Authors:** Bruce M. Boman, Thien-Nam Dinh, Keith Decker, Brooks Emerick, Shirin Modarai, Lynn Opdenaker, Jeremy Z. Fields, Christopher Raymond, Gilberto Schleiniger

## Abstract

The genetic code determines how the precise amino acid sequence of proteins is specified by genomic information in cells. But what specifies the precise histologic organization of cells in plant and animal tissues is unclear. We now hypothesize that another code, the *tissue code*, exists at an even higher level of complexity which determines how tissue organization is dynamically maintained. Accordingly, we modeled spatial and temporal asymmetries of cell division and established that five simple mathematical laws (“the tissue code”) convey a set of biological rules that maintain the specific organization and continuous self-renewal dynamics of cells in tissues. These laws might even help us understand wound healing, and how tissue disorganization leads to birth defects and tissue pathology like cancer.

## INTRODUCTION

Although “the genetic code” explains fidelity of genotypic expression (how nucleotide triplets encode specific amino acids), there is no code as yet that explains the fidelity of tissue organization. For that, one needs to know how cellular mechanisms for tissue renewal encode the organization of cell populations in a tissue, i.e., define a code that explains tissue organization. Like the genetic code, a tissue code would provide a set of rules by which all multicellular organisms maintain themselves in a highly ordered state, but at a higher level of complexity than the genetic code. Scientists assume, it appears, that tissue organization is merely a process of cell proliferation that happens during tissue renewal. But that is insufficient to explain the precision of tissue renewal such as the constancy of tissue size, the subdivision of different tissue types (epithelia, connective tissue, muscle, and nervous tissue), the histologic pattern and unique distribution of specialized cells in each tissue, as well as the precise spatial relationships between individual cells. The tissue code could explain how multicellular tissues are normally organized, how that organization is faithfully maintained through tissue renewal over an organism’s lifetime, and also how healing occurs after injury. Indeed, the existence of plant and animal life on earth for hundreds of million years necessitated an ability to precisely establish and maintain tissue organization.

Determining how the organization of tissues is encoded through tissue renewal presents a highly complex problem in biology. Historically, early efforts toward discovery of the genetic code held that the problem was vastly complex. But, it was mathematics that was instrumental in initial discovery of the genetic code and how the information in DNA is transferred to proteins [1, 2]. Taking this lesson from history, we used mathematical modeling to see if a set of rules could be found that explains how the complex organization of cells in tissues happens in healthy adults. In designing our model, we assumed that asymmetric division is key and must involve temporal as well as spatial mechanisms. An iterative approach was taken to create different models to find a model design that could simulate ongoing maintenance of cellular organization that happens during the continuous turnover of cell populations in a tissue. After many iterations, we identified a set of mathematical laws that may explain how tissues could stay organized. We also then developed computer graphics to visually display the system dynamics.

Another reason that we turned to mathematics was because the mechanisms responsible for maintaining tissue organization were too complex to be easily solved using conventional biological experiments. That would require measurement of the dynamics of different cell types in tissues *in vivo*. Even recent exciting efforts to map whole organisms using image registration and gene expression analysis of all cell types in lower multicellular organisms (e.g. planarians) generate data on only a snapshot in time of the entire animal rather than continual dynamic motion of cells in various tissues [3,4]. Indeed, simultaneous recording of viable cells in terms of their dynamics (rate of cell movement; time of maturation & of division; the positions of cells relative to each other; direction of movement & of division) in living tissues would be necessary to understand the rules for tissue renewal. However, once these rules were identified mathematically, then biological experiments might then be done to validate them.

In previous studies, models for morphogenesis were created to explain how embryonic tissues become organized during development. Indeed, such models have been designed based on mechanics (e.g., adhesion, forces), geometry (e.g. cell shape), pattern formation (e.g. reaction-diffusion), and molecular mechanisms (involving genes, proteins, enzymes, and signaling pathways) [5,6]. And, these models have provided valuable information as to how tissues are spatially formed in the early embryo during development. Some topological models have also been made that describe the geometric spatial organization of adult epithelia [7]. But they do not explain how adult tissue organization is dynamically maintained.

Investigating mechanisms in the dynamics of tissue renewal is a daunting challenge not only for the precision by which tissue organization is unerringly maintained, but also from the rapidity with which tissues replace themselves. In fact, healthy tissue organization in humans is preserved during the replacement of ∼2 trillion cells that turnover every day. This amounts to well over a million, million, million (>10^18^) cell divisions during the ∼60 years of average human adulthood. Given this extensive cell turnover, it is remarkable that the organization of any given tissue is continuously and precisely maintained in our organs. For example, we have been modeling the dynamic properties of tissue renewal of the human colonic epithelium, which takes 3-5 days [8, 9]. During self-renewal in gut, the epithelium stays highly organized in terms of its specific cellular organization as well as distribution and number of crypts. Most other tissue types, even liver and brain, have their characteristic turnover time which continues throughout adulthood [10], as does the precise cellular organization of each tissue. The speed, precision and complexity of tissue renewal processes strongly suggest the existence of an underlying *tissue code*.

To discern rules that define a tissue code, we reasoned that because tissue renewal is integrally linked to maintenance of adult tissue organization, the fidelity of tissue renewal must involve mechanisms that control cell division. We also surmised that any code that maintains tissue organization must contain biological rules that specify: i) precise histologic structure, ii) steady state dynamics, and iii) tissue size. Hence, we conjectured that rules that determine histologic organization involve timing, temporal order, and direction of cell division. We also surmised that once a specific histologic pattern is established, other rules must underlie the steady state dynamics and size of the tissue. Thus, we conjectured that terminal differentiation (whole maturation), cell lifespan, and feedback mechanisms play a role in maintaining steady state dynamics and tissue size. Indeed, it was this line of reasoning that catalyzed our quest to investigate if maintenance of tissue organization can be explained by a set of simple rules.

Other sets of rules have been established (mainly reaction-diffusion models) based on Alan Turing’s ideas, that explain formation of tissue patterns in development. These models typically involve an activator and an inhibitor that diffuses based on an underlying *random* process [11-13]. In contrast, because of the highly organized nature of tissues, in our model, the division and movement of cells was designed to be *non-random* and to give rise to emergent behavior. The classic model that displays emergent behavior is Conway’s game of life [14], but the patterns generated from Conway’s model’s output do not simulate the organization of tissues. In our view, cells within tissues possess the collective intelligence necessary for maintaining tissue organization because they inherit, from parental cells, instructions for timing (duration), temporal order (sequence), and spatial direction of cell division. So, our objective was to build a mathematical model of the tissue code in the form of a set of rules that was general enough and versatile enough that it can be applied to various types of tissue. By specifying settings (i.e. parameter values in our model) for the rules for each tissue type, our tissue code should at least qualitatively, and perhaps even quantitatively, be able to explain the organization of any given tissue type based on the code’s many possible permutations.

Indeed, investigating how normal tissue organization is maintained with high fidelity is becoming one of the most significant questions in science today. For example, one of NSF’s ten big ideas is “Rules of Life”, which provides an agenda for “bold questions that will drive NSF’s long-term research” [15, 16]. To achieve NSF’s goal to “understand the organizational principles and rules of living systems”, scientists will be seeking answers to questions such as how tissues faithfully sustain highly ordered structures, like our quest to seek a code for tissue organization.

## RESULTS

Consequently, we created two mathematical models (one discrete, the other continuous) that account for healthy adult tissue’s main properties: (*i*). Cells are the constituent parts or building blocks that make up tissues. (*ii*). Tissues continuously turnover. (*iii*). The number of cells in tissues is maintained constant. (*iv*). The organization of cell types within tissues is preserved during tissue renewal.

To account for such properties, we designed our model based on temporal asymmetry of cell division as proposed by Spears and Bicknell-Johnson [17]. In this design, division of a mature parent cell produces two progeny cells, a mature (M) cell and an immature (I) cell, with different temporal properties. An earlier version of our model [18], which built upon this temporal asymmetry, simulated dynamics of tissue renewal, but it didn’t explain how adult tissue organization is maintained. Accordingly, we incorporated new mechanisms for timing, temporal order, and spatial direction of cell division (*Rules* 1-5 below). Our discrete model accounts for each cell in the system with rules for spatial and temporal instructions for cell division. Our continuous linear model assumes a large number of cells, and accounts for the relative proportion of each cell type. The continuous model was created to provide quantitative measures of discrete model system dynamics.

### Discrete Model

In creating the discrete model, through extensive iterative modeling, we found only one model design that could simulate tissue organization during the continuous turnover of cell populations. In this design, the maturation period (*c* value) is linked to the degree of rotation (i.e., direction) of cell division, which mimicked the expected emergent behavior of cells in tissues. For this model design, the 5 rules we postulated for tissue organization were as follows:

### *Rule 1*. Timing of cell division

The timing of cell division is based on a fixed cell cycle duration. The timing of cell division of the two different proliferative cell types (mature (M) cells & immature (I) cells) is based on this cell cycle duration.

### *Rule 2*. Temporal order of cell division

Asymmetric cell division of a mature (M) cell generates two cells, the parent M cell and an immature (I) progeny cell, which have different temporal *dynamic* properties. Specifically, the temporal order of division of these two cells (i.e. the sequence in which cell divisions occur over time) is that M cells divide every cell cycle and I cells divide only after a maturation period (*c* value). Once an I cell undergoes maturation, it immediately becomes an M cell and it divides (i.e. undergoing cell division is the condition for maturation).

### *Rule 3*. Spatial direction of cell division

The direction of cell division rotates by a fixed angle, every cycle, which is based on the maturation period (*c* value). During cell division, each M and I daughter cell inherits the instructions for the direction and timing of its next cell division.

### *Rule 4*. Number of cell divisions

The period of time during which M cells can divide to generate I cells is limited by the whole maturation time (*n*_wm_ or age when an M cell becomes wholly mature) which is a function of generation number (*g*). When an M cell reaches this limit, it can no longer proliferate and it becomes a wholly mature (W) cell. Immature (I) cells also become wholly mature if maturation period equals whole maturation time (*c* = *n*_wm_). This rule dictates the number of divisions that a cell will undergo.

### *Rule 5*. Cell lifespan. The lifespan of a cell (*L*) is the time that it exists in the system from the period when it was generated to when it dies

Asymmetric division (*Rule* 2) is modeled as a simple splitting mechanism. Figure 1A shows a cell lineage diagram that illustrates how asymmetric cell division generates an M cell & an I cell, and that I cells divide only after a given maturation period (*c* value). Even in this simple linear case, it is seen that a dynamic self-renewing cell pattern emerges. When rotation of cell division (*Rule* 3) is integrated into this splitting mechanism, more complex self-renewing patterns are generated. In this situation, the *c* value defines the direction that an M cell rotates (degree *R*) during asymmetric division, and the number of leaflets is based on *R*. Figures 1B and 1C show how cellular organization emerges based on the link between maturation (c=6) and rotation (60 degrees). Other settings for *Rule*s 2 & 3 can generate various specific patterns of cells, leaflets, and branches (see Figure S1, Table S1 & agent-based code below).

**Figure 1.**
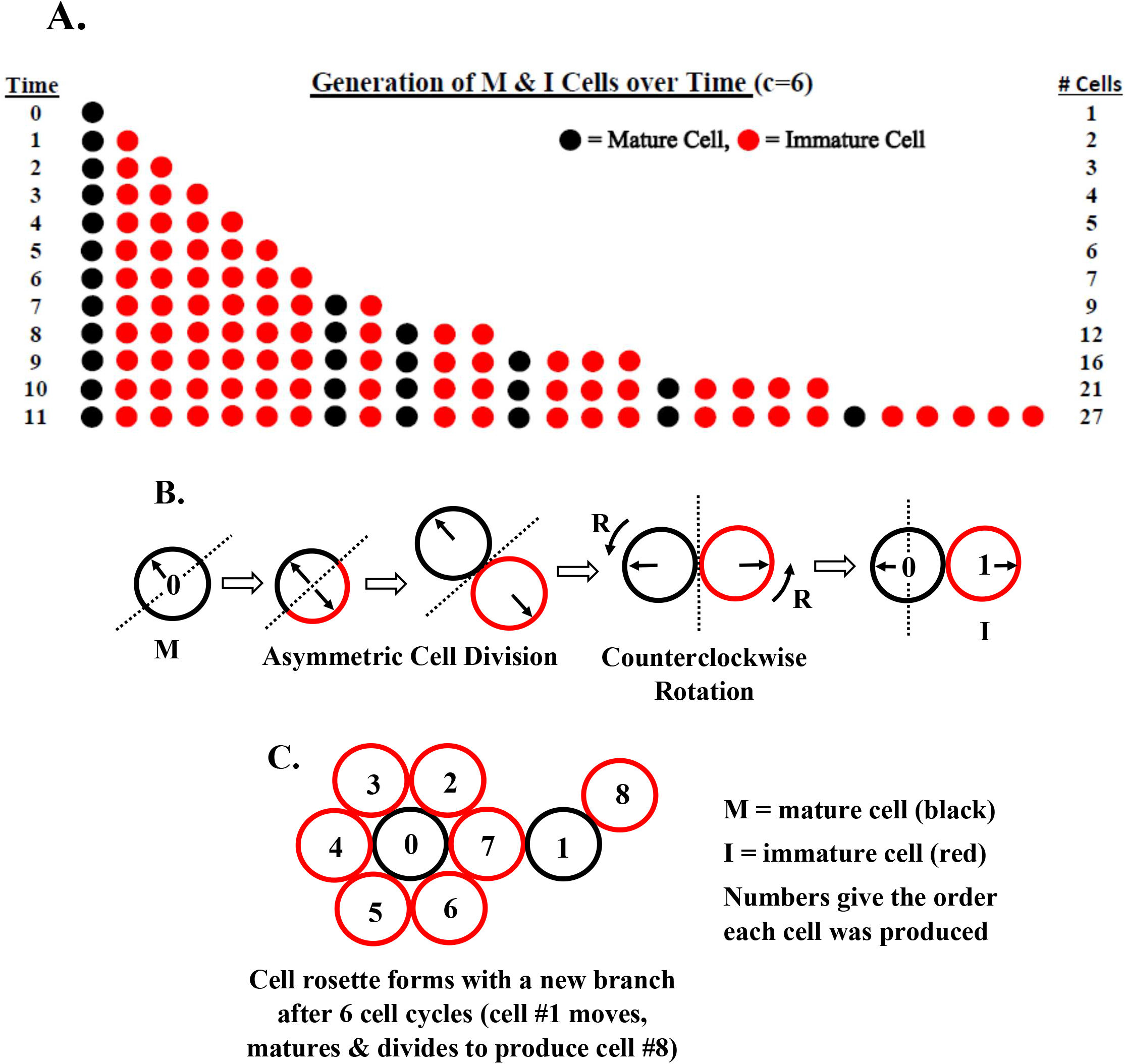
Discrete Model for Asymmetric Cell Division. Panel **A** shows a diagram of the cell lineage production of I & M cells (*Rule 2*) over time for maturation period c=6 based on a simple linear model. Time periods are from top to bottom. Production of cells occurs successively from left to right (new cells displace old cells to the right). It illustrates the process of asymmetric cell division (division of an M cell produces an M cell & an I cell), the time (6 cycles) it takes for I cells to mature into M cells and divide, and the formation of a pattern of cellular organization created by division of M cells within the overall population. Notably, even though all M cells in the population divide every cycle, the cellular organization of the population stays exactly the same. Panels **B & C** show the incorporation of rotation (by 60 degrees) into our model of asymmetric cell division. In other words, instead of I cells being produced to the right of M cells in Panel A, in Panels B and C, it shows how I cells are produced around the M cell. Panel **B** shows modeling according to a simple splitting mechanism (c=6) where cells divide in 60 degree increments determined by line (m) running through the cell center and perpendicular to the M cell (black circle) splitting orientation (solid black arrow). During cell division, both the I cell (red circle) and the M cell (i) inherit a splitting orientation that is the same as the original direction and (ii) rotate by angle *R* counterclockwise about their current orientation (*Rule* 3). The M cell then becomes the grid center and the I cell holds its position relative to the center of the grid. The process repeats itself with the M cell and its new “m” line. When an I cell is produced in an occupied space, the older I cell becomes displaced one position away and in the appropriate direction. Continuation of the process generates an organized pattern of leaflets and branches (Panel **C**). In this way, even though the population of cells is growing, the organization of I and M cells in the population stays exactly the same.

Our discrete model was also formulated as an “agent-based model”. The initial simulations were done based on settings with no differentiation of cells to wholly mature cells (*n*_wm_ = ∞) and cells were immortal (*L* = ∞). For any chosen c value in these runs, the Net Logo code output produces an expanding geometric structure defined by the active division of M cells located within the growing branches that surrounded a clonogenic cell (Figure 2A). While the system in this case does not reach steady-state, the run shows how the model generates symmetric patterns that have a specific organization of cells within branches, and where the organization of cells within the branches remains constant even though the cells within branches are continuously dividing and branches themselves are continually self-renewing and expanding. The two-dimensional structure generated infrom our agent-based model simulates a tissue rosette.

**Figure 2.**
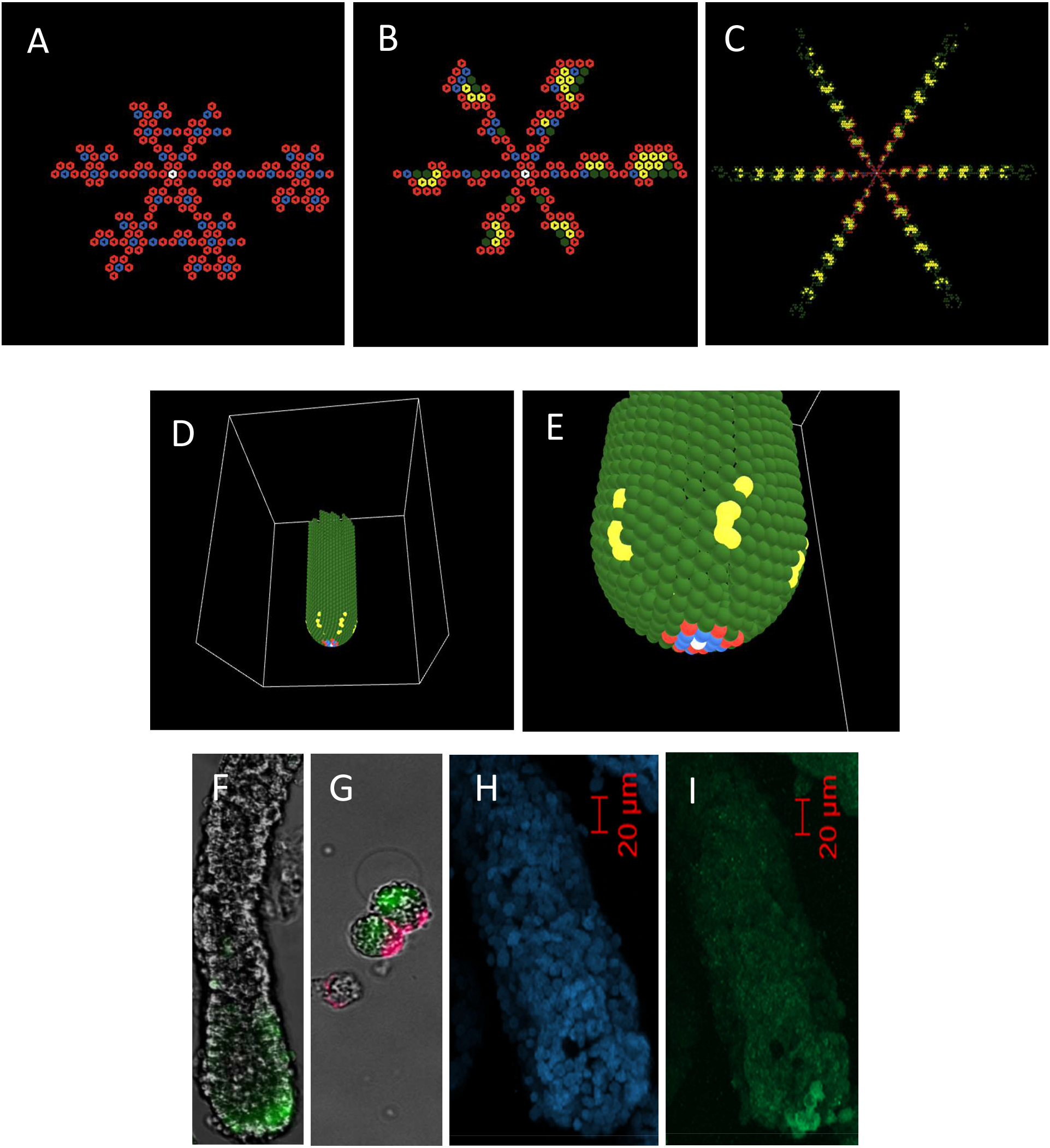
Model output. For any chosen c value in these runs, the computer code output produces an expanding geometric structure defined by the active division of M cells located within the growing branches that surround the clonogenic cell. Panels **A-C** show 2-D output for an agent-based model run based on a maturation time of *c* = 6. Panel **A** shows a snapshot in time of 2D model output (*n*_wm_ = ∞; *L* = ∞ up to time step 24). Panels **B** and **C** show examples of a steady state structure implemented with *c* = 6; *L* = 35, *n*_wm_,_0_ = 11 and *n*_wm_ decreased linearly with *g* (*n*_wm_ = *n*_wm,0_ - *g*). Here, *L* is set large enough to ensure sufficient time for development of the model’s structure while still maintaining steady-state. Panel **B** shows a cell-level view up to time step 27 and Panel **C** up to time step 70. Note the cells eventually die near the edges of the structure. The clonogenic cell is colored white, I cells red, M cells blue, wholly-mature cells green, and overlapping cells yellow. Arrows indicate next split-direction. Panels **D** and **E** show a steady state structure in 3D that simulates organization and dynamics of human colonic crypts (single cell thickness wall with 42 cell columnar circumference). The distribution of various model cell populations (clonogenic, immature, dividing-mature, overlapping, wholly-mature cells) simulates distribution of crypt cell types (stem, progenitor, transit-amplifying, phagocytized apoptotic, terminally-differentiated cells) in biology [8, 9, 53-55, 70-72]. Panels **F-I** show the anatomic location of stem (clonogenic) cells at the bottom of normal human colonic crypts. The location of stem cells in crypts isolated from colonic epithelium was analyzed for the stem cell marker aldehyde dehydrogenase (ALDH) using the ALDEFLUOR assay for ALDH enzymatic activity [55, 73]. Panel **F** shows that ALDH-positive stem cells (green stain) are located at the crypt bottom of viable colonic crypts. Panel **G** reveals an image of live cells from dissociated ALDEFLUOR-stained colonic crypts that were co-stained for the EpCAM epithelial marker (red) to validate that ALDEFLUOR (green) is staining crypt cells not stromal cells. The location of stem cells was also studied using paraformaldehyde-fixed whole colonic crypts. Panel **H** shows a fixed crypt that was stained with DAPI for cell nuclei. Panel **I** shows the same fixed crypt immunostained with anti-ALDH1 antibody revealing that ALDH+ cells reside at the crypt bottom. Thus, model output mimics dynamics of crypt renewal because cells are constantly dividing in regions of active proliferation in the lower crypt while the location and organization of all cell populations and structure (shape, size) stays the same despite a continuous stream of cell populations upward.

To compute the number of cells that are produced in the different branches as a function of time, we developed a generating function (Figure 3). Output on the number of cells generated over time for c = 6 that corresponds to the six branches (Figure 3A) is computed based on this generating function (Figure 3B, C). The numbers generated in growing branches for other c values (c = 2 to 5) are given in Table S2. Model output shows that the total number of cells generated over time (for *n*_wm_ and *L* = ∞) based on any *c* value follows the generalized Fibonacci sequence:

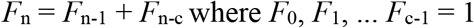

To more closely simulate the dynamic organization of healthy tissues, we then sought to find model settings that establish steady-state tissue renewal. This was done by adjusting *n*_wm_ values while keeping the *c* value constant. An example of a two-dimensional steady state structure generated that simulated a tissue rosette is shown in Figure 2B,C. In establishing a steady state, for any choice of the initial *n*_wm_, a region of active division is generated around the clonogenic cell and expanding branches constantly push out a stream of infertile W cells. Figure 2 shows model output for early time steps (Panel B) and later time steps (Panel C) of the process in which *n*_wm_ is reduced as a function of *g*. While cell lifespan (*L*) does not play a part in the size of the cell production region, it limits the overall size of the structure at steady state conditions. When a steady state structure is projected in three dimensional space (Figure 2D,E), it can simulate the organization and dynamics of tissue structures such as the human colonic crypt (Figure 2F-I).

**Figure 3.**
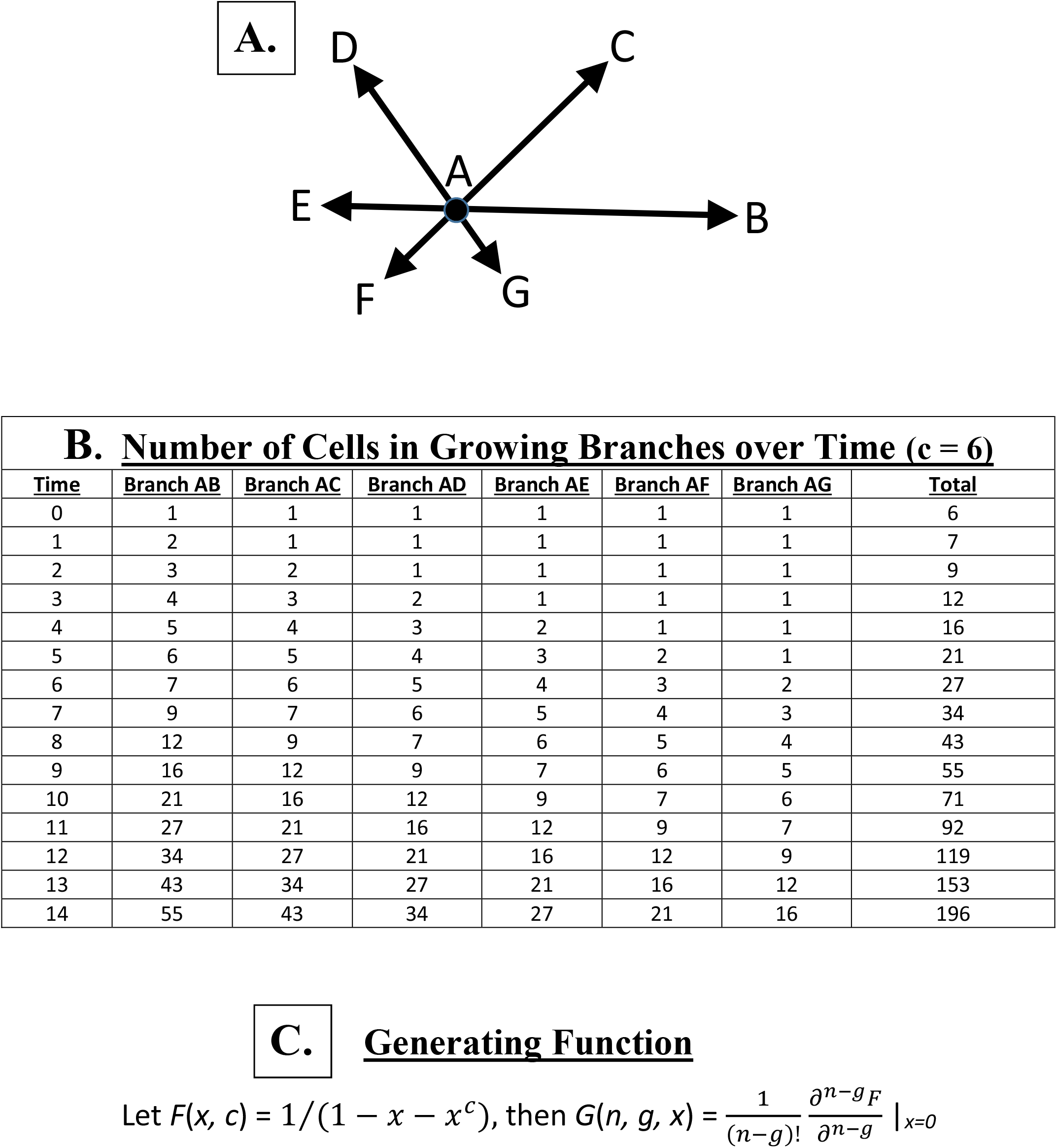
Model design. Our model is designed to produce an expanding geometric structure defined by the active division of M cells located within the growing branches that surround the clonogenic cell (as shown in Fig 2). Panel **A** shows direction of expansion (arrows) of the six branches for a maturation time of *c* = 6. Panel **B** shows output on the number of cells generated over time that corresponds to the six branches (Panel **A**) which is computed based on the generating function (Panel **C**), which gives the number of cells produced in different branches over time for any *c* value. Note that the number of cells for each branch does not include the clonogenic cell or the six cells in the first circumferential layer of leaflet cells surrounding the clonogenic cell. Also note that the number of cells generated over time fits the P-fibonacci sequence for p=6.

Note that the goal of this study was not to create a model for the human colonic crypt, rather our simulation of the crypt and presentation of biological data here was done as a means to show that model output can mimic biological tissue renewal and organization. While a number of important studies on modeling of colonic crypt kinetics have been published [19, 20, reviewed in 21], including our own studies [8, 9], none of the previous models displays emergent cell behavior that simulates the dynamic maintenance of organization of cells in colonic crypts during tissue renewal.

### Continuous Model

In addition to our discrete model, we created a continuous model (Figure 4A) to provide measures of discrete model system dynamics. The rate equations and reaction rates are given in Figure 4B. For the rate constant *k*_1_ for division of M cells, we can choose *k*_1_=1 without loss of generality. This just means that time will be measured in units of 1/*k*_1_ and the other rate constants will therefore be automatically scaled by *k*_1_. The rate constant value (*k*_2_) for division of I cells depends on the *c* value. The W_1 &_ W_2_ cells in the continuous model correspond to W cells in the discrete model. The value of *k*_2_ can be determined by the long-term output on numbers of M and I cells resulting from the discrete model (for each *c* value). The general solution to the system is given in Figure S2, which also provides expressions for the eigenvalues (λ_1,2,3,4_) as follows:

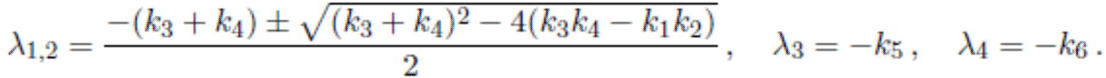

Hence, three eigenvalues are negative (λ_2_ < 0, λ_3_ < 0 and λ_4_ < 0) and the other one, λ_1_ (λ_1_ > λ_2_), is zero, positive or negative depending on whether *k*_3_*k*_4_ = *k*_1_*k*_2_, *k*_3_*k*_4_ < *k*_1_*k*_2_ or *k*_3_*k*_4_ > *k*_1_*k*_2_, respectively. Thus, if *k*_3_*k*_4_ = *k*_1_*k*_2_, then the relative ratios at steady-state are:

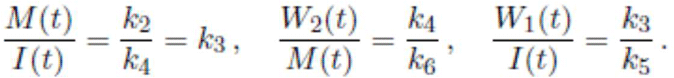

We then studied how the continuous model might provide measures of discrete model system dynamics, particularly of the behavior of cells in the region of active cell division around the clonogenic cell in the discrete model. We did this by comparing the discrete model data with continuous model output when parameter settings in both models produced exponential growth by excluding differentiation of M to W and I to W cells (i.e. *n*_wm_ = ∞; *L* = ∞ in discrete model; *k*_3_ = *k*_4_ = 0 in continuous model). In this case, for the continuous model, we find that 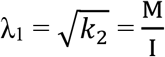(Table 1). For the discrete model, we find that = 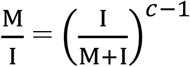 [18]. Solving for 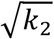 and *c*, the resultant expressions 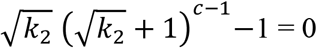 and 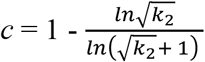 provide a correlation of 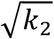 in the continuous model with the *c* value in the discrete model.

**Figure 4.**
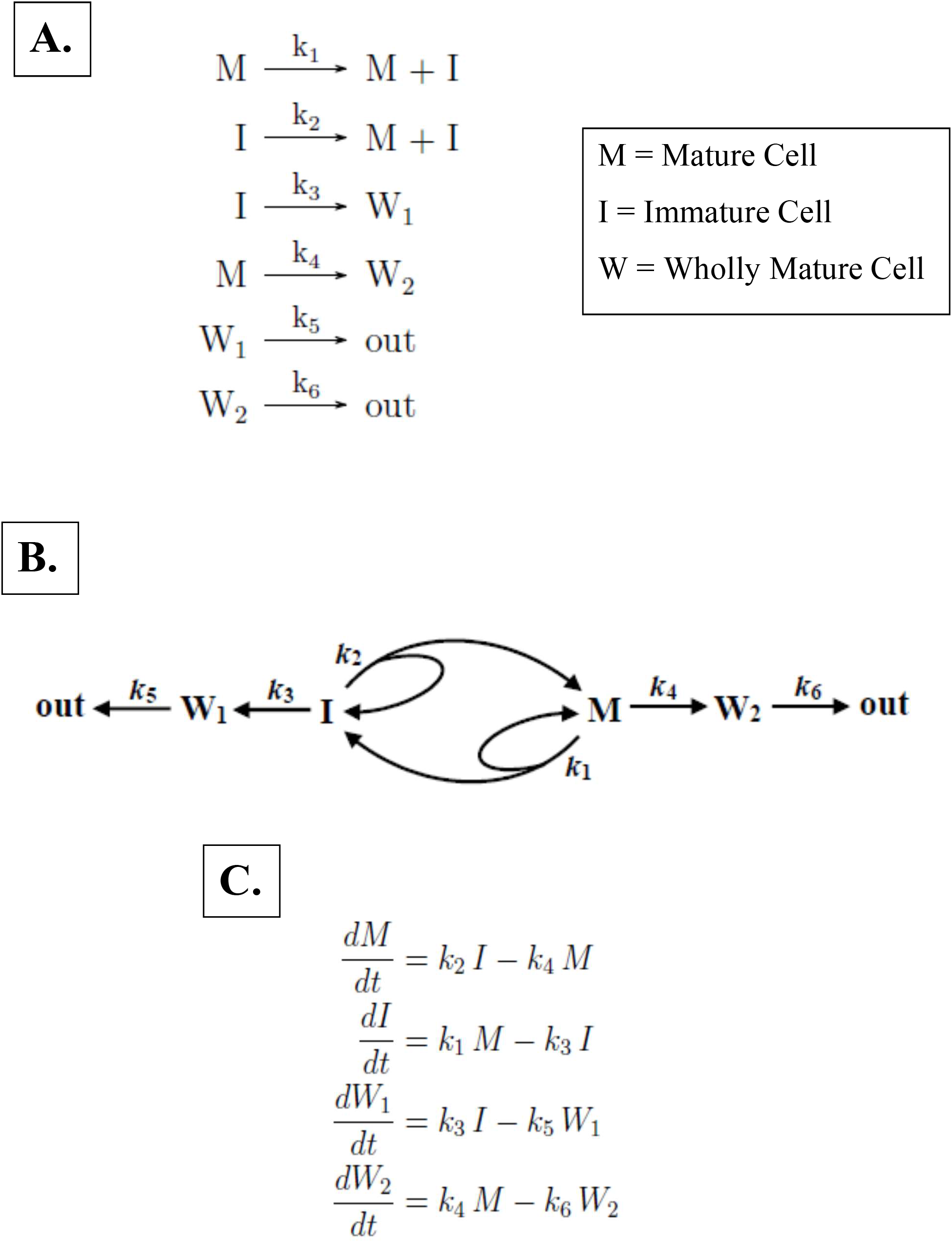
Continuous model for tissue renewal based on asymmetric cell division. Our continuous model has several features that simulate dynamic properties of cells in tissues. The model design is shown in Panel **A**, and rate equations and reaction rates in Panel **B**. The model shows that maturation of I cells and M cells involves their proliferation and differentiation, which occur simultaneously in cells of both populations. Self-renewal occurs on two scales: (i) Individual I and M cells self-renew (governed by *k*_1 &_ *k*_2_ rate constants); (ii) The system itself self-renews. For example, if M cells are depleted, they can be regenerated by division of I cells. Conversely, if I cells are depleted they can be replenished by division of M cells. In other words, there is interconversion between the proliferative cell populations (i.e. between I & M cells), which provides a mechanism for cellular regeneration and, consequently, tissue renewal and even healing after injury. This model design fits with the biology of many tissues, including intestine, lung, stomach and skin, in which proliferative cell populations are known to self-renew as well as to interconvert [74-77]. Asymmetric cell division results in two daughter cells that have different properties. One property is that the two daughter cells exist in different states of maturation which affects their rate of cell division – M cells divide every cycle (based on *k*_1_) whereas I cells divide at a slower rate (based on *k*_2_). An additional property is that the two daughter cells differentiate into different cellular lineages – I cells differentiate into W_1_ cells (based on *k*_3_) and, conversely, M cells differentiate into W_2_ cells (based on *k*_4_). Moreover, based on the model design, it is apparent that different steady states can be achieved in which the proportions of various cell types can differ from one steady state to the next. The achievement of a steady state requires a balance between the production and loss of proliferative cells from the division of M and I cells (governed by *k*_1 &_ *k*_2_) and the whole maturation of W_1_ and W_2_ cells (governed by *k*_3 &_ *k*_4_), respectively. Our modeling also shows, based on eigenvalue solutions, that differentiation along only one lineage will not achieve a steady state – it will only result in exponential growth or decay of the system (data not shown). Thus, differentiation along two lineages in our model is required to achieve a steady state. This result provides an explanation for why many tissues possess two main lineages for differentiation. Examples of such dual lineages in tissues includes hematopoietic (lymphoid & myeloid) [78], brain (neuronal & glial) [79] and intestine (secretory & absorptive) [80] types. Thus, our model design incorporates temporal and spatial properties of cells in tissues including asymmetric division, self-renewal, interconversion, maturation, and steady state kinetics.

**Table 1.** Relationship between rate constant values (k_1_, k_2_), eigenvalue (λ_1_) & ratio of mature (M) to immature (I) cells for different *c* values

| **Table 1. Relationship between rate constant values ($k_1$, $k_2$), eigenvalue ($\lambda_1$) & ratio of mature (M) to immature (I) cells for different $c$ values** |  |  |  |  |
| --- | --- | --- | --- | --- |
| $c$ value | $k_1$ | $k_2$ | $\lambda_1 (= \sqrt{k_2})$ | M/I |
| 1 | 1 | 1 | 1 | 1 |
| 2 | 1 | 0.381966 | 0.618034 | 0.618034 |
| 3 | 1 | 0.216757 | 0.465571 | 0.465571 |
| 4 | 1 | 0.144611 | 0.380278 | 0.380278 |
| 5 | 1 | 0.105442 | 0.324718 | 0.324718 |
| 6 | 1 | 0.081338 | 0.285199 | 0.285199 |

Comparison of model results also revealed that other correlations exist between the two models. For example, discrete model output expressed as a quotient of the sizes of the growing branches for *c* = 2 to *c* = 6 can also be expressed as functions of *k*_2_ rate constant values from the continuous model (Figure S3).

Furthermore, the square roots of the rate constant values for different *c* values relate to the Fibonacci *p*-numbers and golden *p*-sections [22], which indicates that the rate equations and rate constants provide a means to explain growth of the *p*-Fibonacci number recursive series. Overall, results from the continuous model provide measures of discrete system dynamics including temporal evolution of I and M cell numbers and growth of model structures.

## DISCUSSION

The main finding of our modeling of asymmetric cell division is that a simple set of rules based on temporal and spatial rules can generate patterns of cell populations that maintain their organization while continuing to dynamically self-renew, which may explain the fidelity of tissue renewal. How tissue organization is maintained during tissue renewal, to our knowledge, is a question that has not been extensively pursued. One reason might be that in histology, microscopic tissue sections show that tissue organization continually stays constant, making tissues appear static, when in fact tissues are fluid systems, incessantly undergoing renewal.

Another finding from modeling asymmetric cell division is that model output produces cell numbers and organizational patterns that fit with patterns of organization universally found in nature. Specifically, our model output fits the well-known mathematical recursive series – the generalized Fibonacci sequences. Indeed, it is well-known that Fibonacci numbers provide a quantitative description of the phenotype of many living organisms [23-25]. Despite the fact that Fibonacci numbers frequently appear in patterns of growth of plants and animals in nature, the biologic mechanism responsible for the existence of these patterns has not been fully elucidated. The Fibonacci patterns are so common in nature that is seems unlikely that they occur randomly or by chance. Even the geometric proportions of the human body can be described by ratio of Fibonacci numbers (termed the “golden ratio”) [23-25]. Fibonacci number patterns have also been reported to occur at intracellular and cellular scales including the organization of nucleic acid bases in DNA, (termed the “DNA supra code”; [26]), the replication of DNA [27], and clonal growth of human epithelial cells in vitro [28]. This appearance of Fibonacci numbers at different scales – molecular, cellular, organismic – when viewed from the concept of self-similarity suggests that a common underlying mechanism for these Fibonacci patterns exists. In this view, our spatial-temporal asymmetric division mechanism model might lead to an understanding of how Fibonacci numbers arise in patterns of growth of plants and animals and how living organisms are structured at different scales.

Additional key findings from our modeling are: a) daughter cells must inherit instructions for timing, temporal order, and spatial direction of cell division, and b) the direction of cell division is linked to the maturation period. The link between the maturation period and the spatial direction of cell division (*R*) was necessitated by our discovery that, of the numerous model designs we created, the only design that simulated self-renewing tissue organization was the design that incorporated such a link. The production of patterns of cell populations that maintain their organization while continuing to self-renew shows that model output simulates cell dynamics present in biologic tissues. For example, our modeling the dynamics of the human colonic crypt (Figure 2D,E) shows that model output simulates the biology of human crypt organization and structure (Figure 2F-I). Indeed, the growth and maintenance of the model structures – which are generated from model output – occurred due to ongoing cell division within the structure (simulating self-renewal), not just at the ends or edges of the structure (as in crystal growth). Even growth of leaves of plants occurs by cell division throughout the leaf, including within its interior, which gives rise to its specific size and shape. Thus, our results may explain the biology of tissue renewal because in the maintenance of tissues, cells are constantly dividing in regions of active proliferation throughout the tissue while the organization of cells in tissues stays exactly the same.

What biological evidence supports the possibility of a cellular rotational mechanism as proposed in the model? A major feature of our model design is that cells must rotate in a precise coordinated fashion to generate branches that maintain their organization. In biology, there are many known intracellular rotational mechanisms which could serve as the link to the angle of cell division. For example, motor proteins such as ATP synthase rotate 120° [29, 30] and centrosomes and the mitotic spindle rotate at specific angles (usually 90°) during the cell cycle [31-34]. Studies of the motion of cells grown in 2D and 3D cultures also show that cells rotate *in vitro*. When grown on circular 2D substrates (e.g. coated with fibronectin), clusters of cells (from two to several) will coherently rotate together in the absence of any external cues [35-37]. However, when grown in standard adherent 2D tissue culture conditions, cells do not appear to rotate, but they will round up during mitosis and become less firmly attached to the culture substratum. When rounded up, mitotic cells would be less constrained and then have the ability to rotate. When grown in 3D cultures, single cells undergo coordinated rotational movement and even cell clusters up to the four cell stage will continue to cohesively rotate [37-39]. Notably, malignant cells do not display this rotational motion in 3D culture. Mathematical models have been created to explain the mechanics of this rotation of cells grown in 2D and 3D culture conditions [40-42]; but they do not explain, as our model does, how the organization of cells in tissues is maintained during tissue renewal.

What biological evidence supports the idea that the direction of division is linked to the maturation period? Finding the answer to this question involves consideration of two biologic processes: (i) how dividing cells in tissues rotate and (ii) how cells coordinate timing and orientation of cell division. The mechanisms underlying these processes are complex and difficult to quantify because epithelial cells exist in tightly inter-connected epithelial cell sheets *in vivo* that should constrain rotational motility. Still, as early as 1967 it was observed that the orientation of mitotic figures is different in various epithelia [43] and it has now been well-substantiated that orientation of epithelial cells during mitosis is tightly controlled *in vivo*. For example, the orientation of mitosis in the intestine, retina, thyroid and brain is highly precise and regulated in the process called “elevator movement”, which involves the coordination of many cellular mechanisms [44-47]. This process begins with detachment of mitotic cells from the basal lamina in prophase, which gives them the ability to rotate, and the process ends with reattachment of the daughter cells to the basal lamina in late metaphase.

During this process, mitotic cells round up (reminiscent of mitotic cells in 2D cultures) and precisely orient their mitotic spindle, rotate, divide in a specific orientation, and daughter cells re-insert themselves into particular positions within the epithelium. This precise timing and orientation of cell division in biology might begin to be explained by our modeling that cells inherit instructions for timing, temporal order, and spatial direction of cell division. Indeed, inheritance of these instructions was required for the emergent behavior of cells in order for us to explain how the organization of cells in tissues is precisely maintained.

Another key question is: How does model output, which generates rosette structures, fit with similar patterns of cells in nature? It is incorporation of rotation of cell division that generates model rosette structures (see Figures 1,2). This result simulates the arrangement of rosettes found in normal adult tissues, the classic example being rosettes organized as repeating units in the sub-ventricular zone (SVZ) of the brain [48]. Rosettes in the SVZ reside in the neural stem cell niche which is an area of high neurogenic activity that generates new neurons in adults. Even in embryonic stem cell differentiation, the formation of rosette structures is a hallmark of differentiation along the neural lineage [49]. Rosettes are also seen in cancer pathology, particularly brain tumors [50]. And, rosettes are seen in other tissues during embryogenesis [51]. Thus, our model rosette structures that are continuously renewing while in steady state, might fit with the biology of rosettes in the SVZ, which are responsible for tissue renewal in neurogenesis.

While validating that a link exists between the maturation period (*c* value) and the direction of cell division in tissues *in vivo* seems quite challenging, there is some evidence that this link exists. For example, the cell cycle time of enterocytes in the human colonic crypt stem cell niche is long [8,9], and the mitotic spindle of dividing cells aligns perpendicular to the crypt apical surface [52]. In contrast, cell cycle time shortens as proliferative cells migrate up the crypt and the mitotic spindle aligns parallel to the apical surface. Such biological findings indicate the direction of cell division is linked to cell cycle time.

What molecular mechanisms might link these spatial and temporal processes involved in cell division (i.e. maturation period & direction of cell division)? The most logical answer is that it involves a mechanism that links intracellular processes such as large scaffold proteins and hub-type protein complexes that interconnect temporal and spatial processes involved in cell division. For example, the APC and AXIN tumor suppressor proteins are large scaffold proteins that interconnect mitotic spindle orientation and transcriptional regulation, which is crucial to the maintenance of the organization of cell populations in the human colonic crypt [53-57]. Notably, network hub scaffold-type proteins are encoded by many cancer genes, which otherwise, if not mutant, normally function by linking different cellular processes such as proliferation and differentiation [58-60].

Finally, we did this study not only because tissue organization is an important biological phenomenon in it its own right, but also, because identifying mechanisms that cause tissue disorganization might explain tumorigenesis. Specifically, we believed that in order to really understand cancer development, we must understand what controls the organization of normal, healthy tissues. Moreover, elucidation of a tissue code could reveal mechanisms that fill the gap between the pathologists’ and the geneticists’ perspectives on cancer development. From the perspective of some pathologists, tissue disorganization is the root of cancer [61-64]. Indeed, cancer diagnosis universally rests on microscopic evidence of tissue disorganization, invasion and metastasis. From the geneticists’ perspective, on the other hand, cancer is caused by gene mutations. In fact, based on The Cancer Genome Atlas Project, any given cancer carries large numbers of mutations (100s to 1000s) and other molecular alterations [65-67]. Combining these two perspectives, one can argue that tissue disorganization and genetic alterations are both necessary for cancer development, but neither is sufficient to explain cancer. Thus, identification of a tissue code, and understanding how disrupting that code gives rise to tissue pathology, might provide a mechanism that helps explain how gene mutations lead to tissue disorganization and subsequently cancer.

It could also help understand mechanisms that lead to tissue disorganization which contribute to other diseases. Indeed, it is unlikely that tissues can tolerate many errors in their cellular organization without suffering serious impairment of function. One example is disorders of bone structure due to aberrant renewal and remodeling (e.g. osteoporosis). Another example is metaplasia in Barrett’s esophagus. In that disorder, esophageal epithelium becomes changed into a different epithelial type (intestinal) that predisposes to cancer [68]. Indeed, our model predicts that if daughter cells inherit disordered instructions (altered rules), it will lead to tissue disorganization. Therefore, our findings may help us understand how the code for healthy tissue. i.e., the *tissue code*, becomes disrupted in a way that leads to tissue disorganization and to pathology.

## CONCLUSION

In his classic book “What is Life” published years before discovery of DNA and the genetic code, Erwin Schrodinger predicted, based on theoretical reasoning, that chromosomes must contain some kind of a “code-script” that specifies development of an organism. He also stated “The large and important and very much discussed question is: How can the events in space and time which take place within the spatial boundary of a living organism be accounted for by physics and chemistry? [69]. But, even as recent as last year, NSF’s “Rules of Life: Forecasting and Emergence in Living Systems” still states “we do not understand the basic rules that underlie the emergence of multicellular structures” [15, 16].

Our study suggests a possible answer. We discovered, based on modeling of temporal and spatial asymmetries of cell division, that just a few simple rules may explain how living organisms maintain themselves in a highly ordered state despite continuous renewal of their tissues. The “tissue code” could also help understand how the phenotype of an organism emerges through expression of its genes and how tissue pathology (such as cancer) arises from genetic alterations.

Thus, if our hypothesis is biologically validated, it could explain behavior of cells within tissues under normal conditions of persistent tissue renewal, which might catalyze innovation in the field of tissue biology that, unlike histology, may explain how cells in tissues remain *dynamically* organized. It might even provide clues to the evolutionary biologist on how microbes could have evolved into multicellular life 100s of million years ago.

## METHODS (also see Supplemental Material)

### Asymmetric division & rotation

In our basic model, the daughter cell inherits the split-angle of its parent cell (Figure 1B), which turns counterclockwise by one increment every time step. Given this process, leaflets of I cells form around the M cell (Figure 1C). Model output eventually creates branches that emerge from the leaflets.

### Agent-based model

An agent-based model was formulated that incorporates *Rules 2-*5 with the agents being the cells. Maturation (*c*), rotation (*R*), whole-maturation time (*n*_wm_) and lifespan (*L*) specify the age at which cells change state. At age zero, a cell is immature; at age *c*, a cell becomes mature; at age *n*_wm_, a cell becomes wholly-mature; at age *L*, the cell reaches the end of its life cycle and dies. The number of divisions undergone by any given cell is determined by: #divisions = *n*_wm_ − *c*. The clonogenic cell remains immortal because it defines generation zero and continually generates I cells. Thus, its *c* value defines the pattern of the entire structure. The computer code is available upon request.

### Modeling steady-state tissue renewal

To find settings that establish steady-state tissue renewal *n*_*wm*_ values were adjusted while keeping the c value constant such that every M cell under a steady-state condition only produces, on average, a single I cell. Namely, *n*_wm_ was programmed to decrease over time as a function of cell generation (*g*) whereby *n*_wm_ = *n*_wm,0_ − *g*, where *n*_wm,0_ is an initial constant value. Cell generation is the number of divisions removed from the zero^th^ generation clonogenic cell. Since the number of divisions of a cell is defined as *n*_wm_ - *c*, it is clear that when holding *c* constant, *n*_wm_ must be reduced for later generations to reach steady state. Thus, when *n*_wm_ is reduced as a function of *g*, different population sizes will be produced depending on the initial value of *n*_wm_ (*n*_wm,0_).

### Continuous Model

We created a simple, linear continuous model the dynamics of which are defined in terms of rate constant values for division of M and I cells (Figures 4, S2).

### Analysis of Human Colonic Crypts

Colonic crypts were purified from normal human colonic epithelium from surgical specimens as as we previously described [54, 55, 73]. Isolated viable colonic crypts were analyzed for the ALDH stem cell marker by ALDEFLUOR analysis and paraformaldehyde fixed colonic crypts were analyzed for ALDH by imunofluorescence as we previously described [54, 55, 73]. Briefly we used anti-ALDH1 (BD Pharmingen, Franklin Lakes, 1:50) as the primary antibody. The use of human tissues was approved by Institutional Review Board of Christiana Care Health Services, Inc (FWA00006557).

## FOOTNOTES

Author contributions: <u>BM Boman</u>: designed research, performed research, analyzed data, wrote the paper<u>. T-N Dinh</u>: performed research, contributed computational analytic tools, analyzed data, contributed to writing the paper. <u>K Decker</u>: performed research, contributed computational analytic tools, analyzed data. <u>B Emerick</u>: performed research, contributed computational analytic tools, analyzed data. <u>JZ Fields</u>: analyzed data, significantly contributed to writing the paper. C Raymond: analyzed data, contributed to writing the paper. <u>G Schleiniger</u>: designed research, performed research, contributed computational analytic tools, analyzed data, contributed to writing the paper.

## ACKNOWLEDGMENTS

Generous support was provided by the Cancer Code Research Foundation, a UNIDEL Graduate Research Fellowship, and NIH NIGMS Program Grant #P20 GM103446. We thank the following students for their participation in this project: Matt Schmittle, Nathaniel Borders, Ilana Attadgie, Matthew Saponaro, Kris Olli, Scott Fones, Benjamin Clark, Michael Gonzales, Christian Berthia, Raghav Sambasivan, and Megan DiIorio. We thank Dr Nicholas Petrelli for his ongoing support and the late Dr Olaf Runquist for his inspiration and encouragement during our project.

