## Supplemental Methods, Tables 1-3, Figures 1-3 for "Beyond the Genetic Code: *A Tissue Code?*"

### Supplemental Materials

#### **Materials and Methods**.

Experimental Design

We created two mathematical models (one discrete, the other continuous) based on asymmetric cell division. Our discrete model accounts for each cell in the system with rules for timing and direction of cell division and was programed as an agent-based code. Our continuous model assumes a large number of cells, and accounts for the relative proportion of each cell type.

Agent-based model code for the discrete model

In the code, the sum of cell properties and rules defined the analytical description of the model, which was encoded into NetLogo software used to generate output for the actual simulation. In this code, cells have a position that is defined by their place within an implicit tree data structure that is rooted at the clonogenic cell. In this tree, all nodes are cells and all edges represent connections to either its immediate neighbors or empty space. In model runs, the split-angle is continually tracked and updated for I and M cells. The computer code is available upon request from the corresponding author.

Continuous model

Our continuous model for the dynamics of the system is described by rate equations and rate constant values for division of mature and immature cells in the system population. Only the time evolution of the population of each cell type is tracked — no spatial dependence was considered. First-order kinetics was assumed in order to derive a simple linear model that gives insight into the initial stages of cell population growth.

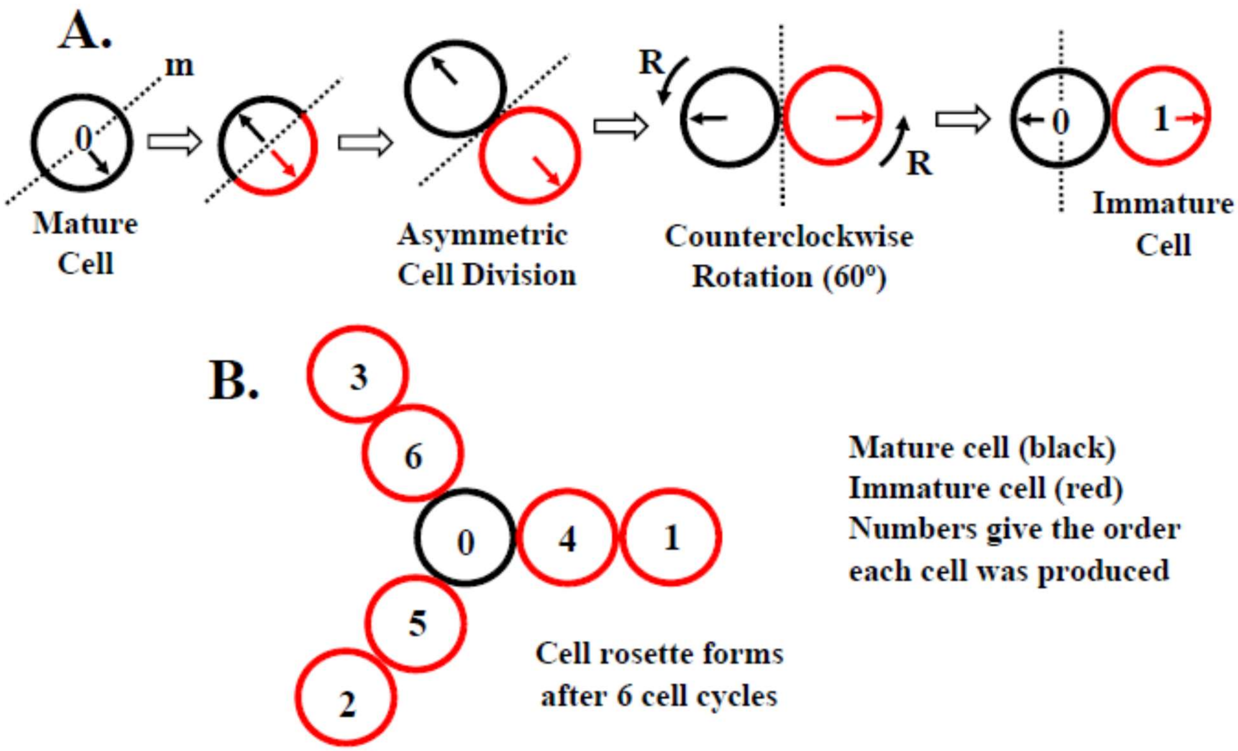


Supplemental Figure 1. Asymmetric division and rotation based on a different set of settings. Panel A shows modeling of cell division based on spatial and temporal asymmetry (Rule 2) according to a simple splitting criterion. Division is modeled as a reflection mechanism determined by a mirror line (m) running through the cell center and perpendicular to the mature cell (black circle) splitting orientation (solid black arrow). After cell division, the mature cell inherits orientation in the opposite direction, while the immature cell (red circle) inherits orientation in the original direction. After cell division, both the mature and immature cells rotate by angle R counterclockwise about their current orientation (Rule 3). After rotation, the mature cell becomes the grid center (i.e., the clonogenic cell) and the immature cell holds its position relative to the center of the grid. Cell division continues with the mature cell and its new mirror line. Cell splitting continues until an immature cell is produced that occupies the space that already contains an immature cell which then becomes displaced one position away and in the appropriate direction. Therefore, the process repeats itself based on the angle of rotation R after cell division and patterns are formed (Panel B).

| $R$ (degrees) | $R$ (rads) | $\ell$ (leaflets) | $N$ (steps) |
|---|---|---|---|
| 180 | $\pi$ | 1 | 3 |
| 0 | 0 | 2 | 5 |
| 60 | $\pi/3$ | 3 | 7 |
| 90 | $\pi/2$ | 4 | 9 |
| 36 | $\pi/5$ | 5 | 11 |
| 120 | $2\pi/3$ | 6 | 13 |
| 45 | $\pi/4$ | 8 | 17 |
| 20 | $\pi/9$ | 9 | 19 |
| 72 | $2\pi/5$ | 10 | 21 |
| 30 | $\pi/6$ | 12 | 25 |
| 12 | $\pi/15$ | 15 | 31 |
| 40 | $2\pi/9$ | 18 | 37 |
| 18 | $\pi/10$ | 20 | 41 |
| 15 | $\pi/12$ | 24 | 49 |
| 24 | $2\pi/15$ | 30 | 61 |
| 10 | $\pi/18$ | 36 | 73 |
| 9 | $\pi/20$ | 40 | 81 |
| 4 | $\pi/45$ | 45 | 91 |
| 6 | $\pi/30$ | 60 | 121 |
| 5 | $\pi/36$ | 72 | 145 |
| 8 | $2\pi/45$ | 90 | 181 |
| 3 | $\pi/60$ | 120 | 241 |
| 2 | $\pi/90$ | 180 | 361 |
| 1 | $\pi/180$ | 360 | 721 |

Supplemental Table 1. The orientation of the mature and immature cells. The orientation of the mature cell at step n, denoted by $\theta_n$, follows the sequence below if it is assumed that the initial orientation of the mature cell is 0 radians or oriented toward the positive x-axis.

$$\theta_1 = 0, \quad \theta_2 = \pi, \quad \theta_n = \theta_{n-2} + R + \pi,$$

Using this relation, the spatial position of the $i_{th}$ daughter cell, denoted $D_i$, can be determined by the following sequence.

$$D_i = \theta_{2i-1} + R, \qquad \text{for } i = 1, 2, \ldots, \frac{N-1}{2}.$$

The $i_{th}$ daughter cell is located on the ray that makes an angle $D_i$ with the x-axis. If the $\theta$ in the previous equation is eliminated, the following formula is obtained.

$$D_i = (i-1)\pi + iR, \qquad \text{for } i = 1, 2, \ldots, \ell,$$

where $\ell$ is the total number of leaflets, $\ell = (N\text{-}1)/2$. Thus, the number of leaflets is related to the given rotational degree, $R$.

**<u>Supplemental Table 2.</u> Generating Function Results for Growing Branch Sizes**

| Generating Function Results for Growing Branch Size (c = 1) | | |
|---|---|---|
| **Time** | **Branch AB** | **Total** |
| 0 | 1 | 1 |
| 1 | 2 | 2 |
| 2 | 4 | 4 |
| 3 | 8 | 8 |
| 4 | 16 | 16 |
| 5 | 32 | 32 |
| 6 | 64 | 64 |
| 7 | 128 | 128 |
| 8 | 256 | 256 |
| 9 | 512 | 512 |
| 10 | 1024 | 1024 |
| 11 | 2048 | 2048 |
| 12 | 4096 | 4096 |
| 13 | 8192 | 8192 |
| 14 | 16384 | 16384 |

| Generating Function Results for Growing Branch Sizes (c = 2) | | | |
|---|---|---|---|
| **Time** | **Branch AB** | **Branch AC** | **Total** |
| 0 | 1 | 1 | 2 |
| 1 | 2 | 1 | 3 |
| 2 | 3 | 2 | 5 |
| 3 | 5 | 3 | 8 |
| 4 | 8 | 5 | 13 |
| 5 | 13 | 8 | 21 |
| 6 | 21 | 13 | 34 |
| 7 | 34 | 21 | 55 |
| 8 | 55 | 34 | 89 |
| 9 | 89 | 55 | 144 |
| 10 | 144 | 89 | 233 |
| 11 | 233 | 144 | 377 |
| 12 | 377 | 233 | 610 |
| 13 | 610 | 377 | 987 |
| 14 | 987 | 610 | 1597 |

| Generating Function Results for Growing Branch Sizes (c = 3) | | | | |
|---|---|---|---|---|
| **Time** | **Branch AB** | **Branch AC** | **Branch AD** | **Total** |
| 0 | 1 | 1 | 1 | 3 |
| 1 | 2 | 1 | 1 | 4 |
| 2 | 3 | 2 | 1 | 6 |
| 3 | 4 | 3 | 2 | 9 |
| 4 | 6 | 4 | 3 | 13 |
| 5 | 9 | 6 | 4 | 19 |
| 6 | 13 | 9 | 6 | 28 |
| 7 | 19 | 13 | 9 | 41 |
| 8 | 28 | 19 | 13 | 60 |
| 9 | 41 | 28 | 19 | 88 |
| 10 | 60 | 41 | 28 | 129 |
| 11 | 88 | 60 | 41 | 189 |
| 12 | 129 | 88 | 60 | 277 |
| 13 | 189 | 129 | 88 | 406 |
| 14 | 277 | 189 | 129 | 595 |

| Generating Function Results for Growing Branch Sizes (c = 4) | | | | | |
|---|---|---|---|---|---|
| **Time** | **Branch AB** | **Branch AC** | **Branch AD** | **Branch AE** | **Total** |
| 0 | 1 | 1 | 1 | 1 | 4 |
| 1 | 2 | 1 | 1 | 1 | 5 |
| 2 | 3 | 2 | 1 | 1 | 7 |
| 3 | 4 | 3 | 2 | 1 | 10 |
| 4 | 5 | 4 | 3 | 2 | 14 |
| 5 | 7 | 5 | 4 | 3 | 19 |
| 6 | 10 | 7 | 5 | 4 | 26 |
| 7 | 14 | 10 | 7 | 5 | 36 |
| 8 | 19 | 14 | 10 | 7 | 50 |
| 9 | 26 | 19 | 14 | 10 | 69 |
| 10 | 36 | 26 | 19 | 14 | 95 |
| 11 | 50 | 36 | 26 | 19 | 131 |
| 12 | 69 | 50 | 36 | 26 | 181 |
| 13 | 95 | 69 | 50 | 36 | 250 |
| 14 | 131 | 95 | 69 | 50 | 345 |

| Generating Function Results for Growing Branch Sizes (c = 5) | | | | | | |
|---|---|---|---|---|---|---|
| **Time** | **Branch AB** | **Branch AC** | **Branch AD** | **Branch AE** | **Branch AF** | **Total** |
| 0 | 1 | 1 | 1 | 1 | 1 | 5 |
| 1 | 2 | 1 | 1 | 1 | 1 | 6 |
| 2 | 3 | 2 | 1 | 1 | 1 | 8 |
| 3 | 4 | 3 | 2 | 1 | 1 | 11 |
| 4 | 5 | 4 | 3 | 2 | 1 | 15 |
| 5 | 6 | 5 | 4 | 3 | 2 | 20 |
| 6 | 8 | 6 | 5 | 4 | 3 | 26 |
| 7 | 11 | 8 | 6 | 5 | 4 | 34 |
| 8 | 15 | 11 | 8 | 6 | 5 | 45 |
| 9 | 20 | 15 | 11 | 8 | 6 | 60 |
| 10 | 26 | 20 | 15 | 11 | 8 | 80 |
| 11 | 34 | 26 | 20 | 15 | 11 | 106 |
| 12 | 45 | 34 | 26 | 20 | 15 | 140 |
| 13 | 60 | 45 | 34 | 26 | 20 | 185 |
| 14 | 80 | 60 | 45 | 34 | 26 | 245 |

**Supplemental Figure 2.** **A simple continuous model**

A simple linear kinetic model of asymmetric cell division is described by the kinetic equations

$$\mathrm{M} \xrightarrow{k_1} \mathrm{M} + \mathrm{I} \qquad\qquad \mathrm{I} \xrightarrow{k_2} \mathrm{M} + \mathrm{I}$$
$$\mathrm{I} \xrightarrow{k_3} \mathrm{W_1} \qquad\qquad \mathrm{M} \xrightarrow{k_4} \mathrm{W_2}$$
$$\mathrm{W_1} \xrightarrow{k_5} \mathrm{out} \qquad\qquad \mathrm{W_2} \xrightarrow{k_6} \mathrm{out}$$

It follows that the differential equations governing the concentrations of mature, immature and wholly-mature cells are

$$\frac{dM}{dt} = k_2\, I - k_4\, M\,, \qquad \frac{dI}{dt} = k_1\, M - k_3\, I \tag{1}$$
$$\frac{dW_1}{dt} = k_3\, I - k_5\, W_1\,, \qquad \frac{dW_2}{dt} = k_4\, M - k_6\, W_2 \tag{2}$$

Let $\mathbf{c}$ be the concentration vector: $\mathbf{c} = (M, I, W_1, W_2)$. The equivalent system is

$$\frac{d\mathbf{c}}{dt} = A\,\mathbf{c}\,, \quad \text{where} \quad A = \begin{bmatrix} -k_4 & k_2 & 0 & 0 \\ k_1 & -k_3 & 0 & 0 \\ 0 & k_3 & -k_5 & 0 \\ k_4 & 0 & 0 & -k_6 \end{bmatrix}, \tag{3}$$

and the characteristic equation is

$$\left[\lambda^2 + (k_3 + k_4)\lambda + k_3 k_4 - k_1 k_2\right](\lambda + k_5)(\lambda + k_6) = 0.$$

The eigenvalues are

$$\lambda_{1,2} = \frac{-(k_3 + k_4) \pm \sqrt{(k_3 + k_4)^2 - 4(k_3 k_4 - k_1 k_2)}}{2}\,, \quad \lambda_3 = -k_5\,, \quad \lambda_4 = -k_6\,.$$

Hence, three eigenvalues are negative ($\lambda_2 < 0$, $\lambda_3 < 0$ and $\lambda_4 < 0$) and the other, $\lambda_1$ ($\lambda_1 > \lambda_2$), is zero, positive or negative depending on whether $k_3 k_4 = k_1 k_2$, $k_3 k_4 < k_1 k_2$ or $k_3 k_4 > k_1 k_2$, respectively. In order to avoid exponential growth, $k_3 k_4 \geq k_1 k_2$ must hold.

The solution can be easily obtained in terms of the initial values of the cell populations, $M_0$, $I_0$, $W_{1,0}$ and $W_{2,0}$. Let

$$K = \sqrt{(k_3 + k_4)^2 - 4(k_3 k_4 - k_1 k_2)} = \sqrt{(k_3 - k_4)^2 + 4 k_1 k_2}\,.$$

The general solution of (1),(2) is

$$M(t) = 2k_2 \left(c_1\, e^{\lambda_1 t} + c_2\, e^{\lambda_2 t}\right), \quad I(t) = c_1\,(k_4 - k_3 + K)\, e^{\lambda_1 t} + c_2\,(k_4 - k_3 - K)\, e^{\lambda_2 t}, \tag{4}$$

$$W_1(t) = k_3 \left(c_1\, \frac{k_4 - k_3 + K}{\lambda_1 + k_5}\, e^{\lambda_1 t} + c_2\, \frac{k_4 - k_3 - K}{\lambda_2 + k_5}\, e^{\lambda_2 t}\right) + c_3\, e^{\lambda_3 t}, \tag{5}$$

$$W_2(t) = 2k_2 k_4 \left(\frac{c_1}{\lambda_1 + k_6}\, e^{\lambda_1 t} + \frac{c_2}{\lambda_2 + k_6}\, e^{\lambda_2 t}\right) + c_4\, e^{\lambda_4 t}, \tag{6}$$

where the constants $c_1$ , $c_2$ , $c_3$ and $c_4$ are determined from the initial populations. The expressions (4)–(6) assume that $\lambda_1$ and $\lambda_2$ are distinct from both $\lambda_3$ and $\lambda_4$ . If this is not the case, similar expressions are obtained and the behavior discussed below is the same.

It is convenient to non-dimensionalize time by $1/k_1$ , the characteristic time for mature cell division, i.e. time will be measured in units of $1/k_1$. This is equivalent to replacing $k_j$ by $k_j/k_1$ in equation (1),(2), or setting $k_1 = 1$ throughout.

In all cases, the ratio of mature to immature cells in the long run is

$$\frac{M(t)}{I(t)} \approx \frac{2k_2}{K + k_4 - k_3} = \frac{2k_2}{k_4 - k_3 + \sqrt{(k_4 - k_3)^2 + 4k_2}} \,. \tag{7}$$

The constants in (4)–(6) are

$$c_1 = \frac{2k_2 I_0 + (k_3 - k_4 + K)\, M_0}{4k_2 K} \,, \quad c_2 = \frac{(k_4 - k_3 + K)\, M_0 - 2k_2 I_0}{4k_2 K} \,, \tag{8}$$

$$c_3 = W_{1,0} - \frac{k_3}{2K} \left[ \frac{(K + k_4 - k_3) I_0 + 2M_0}{\lambda_1 + k_5} + \frac{(K + k_3 - k_4) I_0 - 2M_0}{\lambda_2 + k_5} \right] , \tag{9}$$

$$c_4 = W_{2,0} - \frac{k_4}{2K} \left[ \frac{2k_2 I_0 + (k_3 - k_4 + K)\, M_0}{\lambda_1 + k_6} + \frac{(k_4 - k_3 + K)\, M_0 - 2k_2 I_0}{\lambda_2 + k_6} \right] . \tag{10}$$

The limiting relative ratios (for large $t$) are

$$\frac{M(t)}{I(t)} \approx \frac{2k_2}{k_4 - k_3 + K} \,, \quad \frac{W_2(t)}{M(t)} \approx \frac{2k_4}{2k_6 - k_3 - k_4 + K} \,, \quad \frac{W_1(t)}{I(t)} \approx \frac{2k_3}{2k_5 - k_3 - k_4 + K} \,.$$

A nonzero steady-state for this simple model can only exist if $\lambda_1 = 0$, i.e. only if $k_3 k_4 = k_2$ , in which case $K = k_3 + k_4$ , $\lambda_2 = -(k_3 + k_4)$ and the relative ratios at steady-state are

$$\frac{M(t)}{I(t)} = \frac{k_2}{k_4} = k_3 \,, \quad \frac{W_2(t)}{M(t)} = \frac{k_4}{k_6} \,, \quad \frac{W_1(t)}{I(t)} = \frac{k_3}{k_5} \,.$$

The unique, asymptotically stable, steady-state is given by

$$M(t) = \frac{k_2 I_0 + k_3 M_0}{k_3 + k_4} \,, \quad I(t) = \frac{k_2 I_0 + k_3 M_0}{k_3 (k_3 + k_4)} \,, \tag{11}$$

$$W_1(t) = \frac{k_2 I_0 + k_3 M_0}{k_5 (k_3 + k_4)} \,, \quad W_2(t) = \frac{k_4 (k_2 I_0 + k_3 M_0)}{k_6 (k_3 + k_4)} \,. \tag{12}$$

Modeling the initial stage of tissue formation, prior to any cell reaching whole maturation, is equivalent to setting $k_3 = k_4 = 0$ in (1),(2), and $W_{1,0} = W_{2,0} = 0$, in which case $\lambda_{1,2} = \pm\sqrt{k_2}$ , $K = 2\sqrt{k_2}$ , and from (8)–(10)

$$c_1 = \frac{1}{4\sqrt{k_2}} \left( \frac{M_0}{\sqrt{k_2}} + I_0 \right) , \quad c_2 = \frac{1}{4\sqrt{k_2}} \left( \frac{M_0}{\sqrt{k_2}} - I_0 \right) , \quad c_3 = c_4 = 0 \,.$$

Hence, in the initial stages of tissue formation, from (4) we obtain

$$M(t) = \sqrt{k_2}\,\frac{e^{\sqrt{k_2}t}}{2}\left[\frac{M_0}{\sqrt{k_2}} + I_0 + \left(\frac{M_0}{\sqrt{k_2}} - I_0\right)e^{-2\sqrt{k_2}t}\right] \tag{13}$$

$$I(t) = \frac{e^{\sqrt{k_2}t}}{2}\left[\frac{M_0}{\sqrt{k_2}} + I_0 - \left(\frac{M_0}{\sqrt{k_2}} - I_0\right)e^{-2\sqrt{k_2}t}\right] \tag{14}$$

so that, even for a relatively short time (in units of characteristic time of mature cell division),

$$\frac{M(t)}{I(t)} \approx \sqrt{k_2}\,.$$

Therefore, consistency between the continuous model and the discrete model requires that this relation holds for the latest times of the initial stage of tissue formation. This is consistent with the idea proposed in the Introduction of a set of biologic rules that encodes information in cells for histologic structure similar to the information contained in the meristem of a plant that determines its specific structure and organization. The initial stages of tissue development are determined by the rates of mature and immature cell division, $k_1$ and $k_2$, where $k_1 = 1$ follows from choosing the time scale to be $1/k_1$, and the value of $k_2$ follows from the rules described in the discrete model.

Supplemental Figure 3: Relationship between discrete model output expressed as a quotient of growing branch sizes for $c = 2$ to $c = 6$ and the $k_2$ rate constant values from the continuous model.

**c = 2**

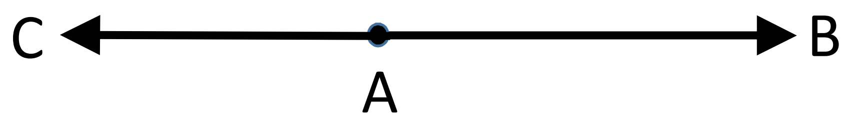


| System Relationships | Correlation with $k_2$ Rate Constant | Value |
|---|---|---|
| $\frac{AC}{AB + AC}$ | $k_2$ | 0.381966 |
| $\frac{AB + AC}{AC}$ | $\frac{1}{k_2} = 2+\sqrt[2]{k_2}$ | 2.618034 |
| $\frac{AC}{AB} = \frac{AB}{CB} = \frac{AB}{AB+AC}$ | $\sqrt[2]{k_2} = 1\text{-}k_2$ | 0.618034 |
| $\frac{AB}{AC} = \frac{CB}{AB} = \frac{AB+AC}{AB}$ | $\frac{1}{\sqrt[2]{k_2}} = 1+\sqrt[2]{k_2}$ | 1.618034 |

| Polynomial Expressions & Root Values* | | |
|---|---|---|
| $k_2$ | $x^2 - 3x + 1 = 0$ | $X_1 = 2.618034$<br>$X_2 = 0.381967$ |
| $\frac{1}{k_2} = 2+\sqrt[2]{k_2}$ | $x^2 - 3x + 1 = 0$ | $X_1 = 2.618034$<br>$X_2 = 0.381967$ |
| $\sqrt[2]{k_2} = 1\text{-}k_2$ | $x^2 + x - 1 = 0$ | $X_1 = 0.618034$<br>$X_2 = -1.618034$ |
| $\frac{1}{\sqrt[2]{k_2}} = 1+\sqrt[2]{k_2}$ | $x^2 - x - 1 = 0$ | $X_1 = 1.618034$<br>$X_2 = -0.618034$ |

*See Table S3 for exact root values

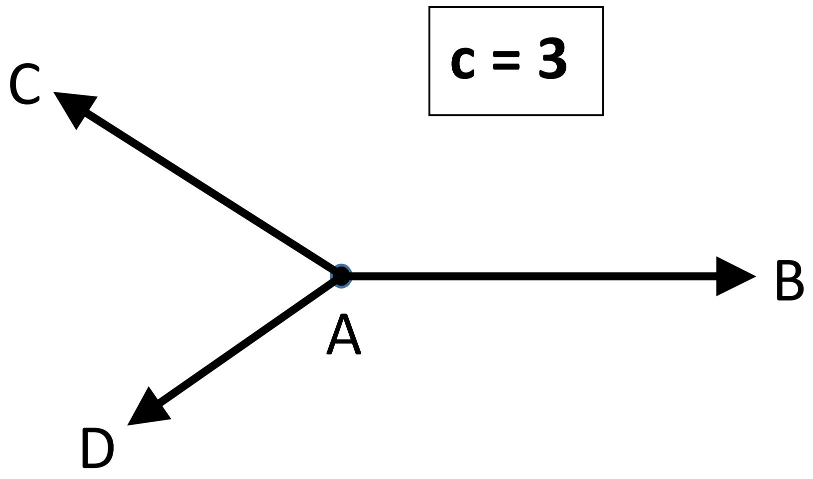


| System Relationships | Correlation with $k_2$ Rate Constant | Value |
|---|---|---|
| $\frac{AD}{AB + AC + AD}$ | $k_2$ | 0.216757 |
| $\frac{AD}{AB} = \frac{AC}{AB+AD} = \frac{AB}{AB+AC+AD}$ | $\sqrt[2]{k_2}$ | 0.465571 |
| $\frac{AB}{AD} = \frac{AB+AC+AD}{AB}$ | $\frac{1}{\sqrt[2]{k_2}}$ | 2.147900 |
| $\frac{AC}{AB} = \frac{AD}{AC} = \frac{AB}{AB+AD}$ | $\sqrt[4]{k_2}$ | 0.682328 |
| $\frac{AB}{AC} = \frac{AC}{AD} = \frac{AB+AD}{AB}$ | $\frac{1}{\sqrt[4]{k_2}} = 1 + \sqrt[2]{k_2}$ | 1.465571 |
| $\frac{AC}{AB+AC+AD} = \frac{AD}{AB+AD}$ | $1 - \sqrt[4]{k_2}$ | 0.317672 |
| $\frac{AB+AC+AD}{AC} = \frac{AB+AD}{AD}$ | $\frac{1}{1 - \sqrt[4]{k_2}} = 1 + \frac{1}{\sqrt[2]{k_2}}$ | 3.1479 |

| Polynomial Expressions & Root Values* | | |
|---|---|---|
| $k_2$ | $x^3 - 2x^2 + 5x - 1 = 0$ | $X_1 = 0.21676$<br>$X_2 = 0.89162+1.95409*i$<br>$X_3 = 0.89162-1.95409*i$ |
| $\sqrt[2]{k_2}$ | $x^3 + 2x^2 + x - 1 = 0$ | $X_1 = 0.46557$<br>$X_2 = -1.23279+0.79255*i$<br>$X_3 = -1.23279-0.79255*i$ |
| $\frac{1}{\sqrt[2]{k_2}}$ | $x^3 - x^2 - 2x - 1 = 0$ | $X_1 = 2.1479$<br>$X_2 = -0.57395+0.36899*i$<br>$X_3 = -0.57395-0.36899*i$ |
| $\sqrt[4]{k_2}$ | $x^3 + x - 1 = 0$ | $X_1 = 0.68233$<br>$X_2 = -0.34116+1.16154*i$<br>$X_3 = -0.34116-1.16154*i$ |
| $\frac{1}{\sqrt[4]{k_2}} = 1 + \sqrt[2]{k_2}$ | $x^3 - x^2 - 1 = 0$ | $X_1 = 1.46557$<br>$X_2 = -0.23279+0.79255*i$<br>$X_3 = -0.23279-0.79255*i$ |
| $1 - \sqrt[4]{k_2}$ | $x^3 - 3x^2 + 4x - 1 = 0$ | $X_1 = 0.31767$<br>$X_2 = 1.34116+1.16154*i$<br>$X_3 = 1.34116-1.16154*i$ |
| $\frac{1}{1 - \sqrt[4]{k_2}} = 1 + \frac{1}{\sqrt[2]{k_2}}$ | $x^3 - 4x^2 + 3x - 1 = 0$ | $X_1 = 3.1479$<br>$X_2 = 0.42605+0.36899*i$<br>$X_3 = 0.42605-0.36899*i$ |

*See Table S3 for exact root values

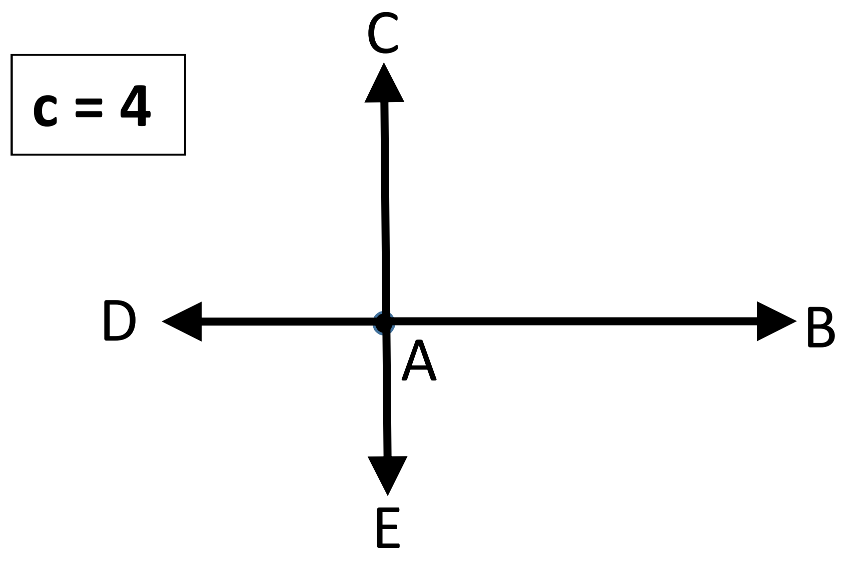


| System Relationships | Correlation with $k_2$ Rate Constant | Value |
|---|---|---|
| $\frac{AE}{AB + AC + AD + AE}$ | $k_2$ | 0.144611 |
| $\frac{AE}{AB} = \frac{AB}{AB+AC+AD+AE} = \frac{AC}{AB+AD+AE} = \frac{AD}{AB+AE}$ | $\sqrt[2]{k_2}$ | 0.380277 |
| $\frac{AB}{AE} = \frac{AB+AC+AD+AE}{AB} = \frac{AB+AD+AE}{AC} = \frac{AB+AE}{AD}$ | $\frac{1}{\sqrt[2]{k_2}}$ | 2.629662 |
| $\frac{AC}{AB} = \frac{AD}{AC} = \frac{AE}{AD} = \frac{AB}{AB+AE}$ | $\sqrt[6]{k_2}$ | 0.724491 |
| $\frac{AB}{AC} = \frac{AC}{AD} = \frac{AD}{AE} = \frac{AB+AE}{AB}$ | $\frac{1}{\sqrt[6]{k_2}} = 1 + \sqrt[2]{k_2}$ | 1.380277 |

| | | |
|---|---|---|
| $\frac{AC}{AB+AC+AD+AE} = \frac{AD}{AB+AD+AE} = \frac{AE}{AB+AE}$ | $1 - \sqrt[6]{k_2}$ | 0.275508 |
| $\frac{AB+AC+AD+AE}{AC} = \frac{AB+AD+AE}{AD} = \frac{AB+AE}{AE}$ | $\frac{1}{1-\sqrt[6]{k_2}} = 1+\frac{1}{\sqrt[2]{k_2}}$ | 3.629659 |
| $\frac{AD}{AB} = \frac{AE}{AC}$ | $\sqrt[3]{k_2}$ | 0.524888 |
| $\frac{AB}{AD} = \frac{AC}{AE}$ | $\frac{1}{\sqrt[3]{k_2}}$ | 1.905168 |

| Polynomial Expressions & Root Values* for c = 4 | | |
|---|---|---|
| $k_2$ | $x^4 - 3x^3 + x^2 - 7x + 1 = 0$ | **$X_1$ = 0.144611**<br>**$X_2$ = 3.30939**<br>**$X_3$ = -0.227+1.42759*i**<br>**$X_4$ = -0.227-1.42759*i** |
| $\sqrt[2]{k_2}$ | $x^4 + 3x^3 + 3x^2 + x - 1 = 0$ | **$X_1$ = 0.380277**<br>**$X_2$ = -1.81917**<br>**$X_3$ = -0.78055+0.91447*i**<br>**$X_4$ = -0.78055-0.91447*i** |
| $\frac{1}{\sqrt[2]{k_2}}$ | $x^4 - x^3 - 3x^2 - 3x - 1 = 0$ | **$X_1$ = -0.5497**<br>**$X_2$ = 2.62966**<br>**$X_3$ = -0.53998+0.63262*i**<br>**$X_4$ = -0.53998-0.63262*i** |

| | | |
|---|---|---|
| $\sqrt[6]{k_2}$ | $x^4 + x - 1 = 0$ | **$X_1$ = 0.724491**<br>**$X_2$ = -1.22074**<br>**$X_3$ = 0.24813+1.03398*i**<br>**$X_4$ = 0.24813-1.03398*i** |
| $\frac{1}{\sqrt[6]{k_2}} = 1 + \sqrt[2]{k_2}$ | $x^4 - x^3 - 1 = 0$ | **$X_1$ = -0.81917**<br>**$X_2$ = 1.38028**<br>**$X_3$ = 0.21945+0.91447*i**<br>**$X_4$ = 0.21945-0.91447*i** |
| $1 - \sqrt[6]{k_2}$ | $x^4 - 4x^3 + 6x^2 - 5x + 1 = 0$ | **$X_1$ = 0.275508**<br>**$X_2$ = 2.22074**<br>**$X_3$ = 0.75187+1.03398*i**<br>**$X_4$ = 0.75187-1.03398*i** |
| $\frac{1}{1 - \sqrt[6]{k_2}} = 1 + \frac{1}{\sqrt[2]{k_2}}$ | $x^4 - 5x^3 + 6x^2 - 4x + 1 = 0$ | **$X_1$ = 0.4503**<br>**$X_2$ = 3.62966**<br>**$X_3$ = 0.46002+0.63262*i**<br>**$X_4$ = 0.46002-0.63262*i** |
| $\sqrt[3]{k_2}$ | $x^4 - 2x^2 - x + 1 = 0$ | **$X_1$ = 0.524888**<br>**$X_2$ = 1.49022**<br>**$X_3$ = -1.00755+0.51312*i**<br>**$X_4$ = -1.00755-0.51312*i** |

| | | |
|---|---|---|
| $\frac{1}{\sqrt[3]{k_2}}$ | $x^4 - x^3 - 2x^2 + 1 = 0$ | **$X_1$ = 0.67104**<br>**$X_2$ = 1.90517**<br>**$X_3$ = -0.7881+0.40136*i**<br>**$X_4$ = -0.7881-0.40136*i** |

*Exact expressions for the roots of the fourth degree polynomials are available, but their long and complicated expressions are not of special interest for this work.

**c = 5**

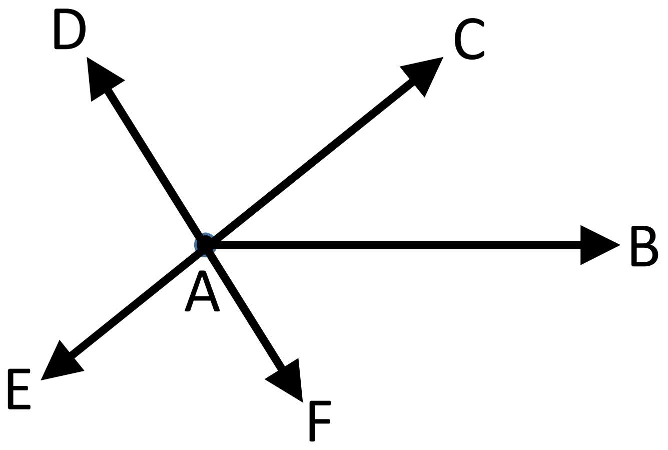


| System Relationships | Correlation with $k_2$ Rate Constant | Value |
|---|---|---|
| $\frac{AF}{AB + AC + AD + AE + AF}$ | $k_2$ | 0.1054418 |
| $\frac{AB}{AB+AC+AD+AE+AF} = \frac{AC}{AB+AD+AE+AF} = \frac{AD}{AB+AE+AF} = \frac{AE}{AB+AF} = \frac{AF}{AB}$ | $\sqrt[2]{k_2}$ | 0.324718 |
| $\frac{AB+AC+AD+AE+AF}{AB} = \frac{AB+AD+AE+AF}{AC} = \frac{AB+AE+AF}{AD} = \frac{AB+AF}{AE} = \frac{AB}{AF}$ | $\frac{1}{\sqrt[2]{k_2}}$ | 3.079595 |
| $\frac{AC}{AB} = \frac{AD}{AC} = \frac{AE}{AD} = \frac{AF}{AE} = \frac{AB}{AB+AF}$ | $\sqrt[8]{k_2}$ | 0.754878 |
| $\frac{AB}{AC} = \frac{AC}{AD} = \frac{AD}{AE} = \frac{AE}{AF} = \frac{AB+AF}{AB}$ | $\frac{1}{\sqrt[8]{k_2}} = 1 + \sqrt[2]{k_2}$ | 1.324718 |

| | | |
|---|---|---|
| $\frac{AC}{AB+AC+AD+AE+AF} = \frac{AD}{AB+AD+AE+AF} = \frac{AE}{AB+AE+AF} = \frac{AF}{AB+AF}$ | $1 - \sqrt[8]{k_2}$ | 0.245122 |
| $\frac{AB+AC+AD+AE+AF}{AC} = \frac{AB+AD+AE+AF}{AD} = \frac{AB+AE+AF}{AE} = \frac{AB+AF}{AF}$ | $\frac{1}{1-\sqrt[8]{k_2}} = 1 + \frac{1}{\sqrt[2]{k_2}}$ | 4.07960 |
| $\frac{AD}{AB} = \frac{AE}{AC} = \frac{AF}{AD}$ | $\sqrt[4]{k_2} = \frac{1}{1+\sqrt[8]{k_2}}$ | 0.569840 |
| $\frac{AB}{AD} = \frac{AC}{AE} = \frac{AD}{AF}$ | $\frac{1}{\sqrt[4]{k_2}} = 1 + \sqrt[8]{k_2}$ | 1.754878 |

| Polynomial Expressions & Root Values* | | |
|---|---|---|
| $k_2$ | $x^5 - 4x^4 + 6x^3 + 4x^2 + 9x - 1 = 0$ | **$X_1$ = 0.10544**<br>**$X_2$ = -0.5+0.86603*i**<br>**$X_3$ = -0.5-0.86603*i**<br>**$X_4$ = 2.44728+1.86942*i**<br>**$X_5$ = 2.44728-1.86942*i** |
| $\sqrt[2]{k_2}$ | $x^5 + 4x^4 + 6x^3 + 4x^2 + x - 1 = 0$ | **$X_1$ = 0.32472**<br>**$X_2$ = -0.5+0.86603*i**<br>**$X_3$ = -0.5-0.86603*i**<br>**$X_4$ = -1.66236+0.56228*i**<br>**$X_5$ = -1.66236-0.56228*i** |
| $\frac{1}{\sqrt[2]{k_2}}$ | $x^5 - x^4 - 4x^3 - 6x^2 - 4x - 1 = 0$ | **$X_1$ = 3.0796**<br>**$X_2$ = -0.5398+0.18258*i** |

| | | |
|---|---|---|
| | | $X_3$ = -0.5398-0.18258*i<br>$X_4$ = -0.5+0.86603*i<br>$X_5$ = -0.5-0.86603*i |
| $\sqrt[8]{k_2}$ | $x^5 + x - 1 = 0$ | $X_1$ = 0.75488<br>$X_2$ = 0.5+0.86603*i<br>$X_3$ = 0.5-0.86603*i<br>$X_4$ = -0.87744+0.74486*i<br>$X_5$ = -0.87744-0.74486*i |
| $\frac{1}{\sqrt[8]{k_2}} = 1 + \sqrt[2]{k_2}$ | $x^5 - x^4 - 1 = 0$ | $X_1$ = 1.32472<br>$X_2$ = 0.5+0.86603*i<br>$X_3$ = 0.5-0.86603*i<br>$X_4$ = -0.66236+0.56228*i<br>$X_5$ = -0.66236-0.56228*i |
| $1 - \sqrt[8]{k_2}$ | $x^5 - 5x^4 + 10x^3 - 10x^2 + 6x - 1 = 0$ | $X_1$ = 0.24512<br>$X_2$ = 0.5+0.86603*i<br>$X_3$ = 0.5-0.86603*i<br>$X_4$ = 1.87744+0.74486*i<br>$X_5$ = 1.87744-0.74486*i |
| $\frac{1}{1 - \sqrt[8]{k_2}} = 1 + \frac{1}{\sqrt[2]{k_2}}$ | $x^5 - 6x^4 + 10x^3 - 10x^2 + 5x - 1 = 0$ | $X_1$ = 4.0796<br>$X_2$ = 0.4602+0.18258*i<br>$X_3$ = 0.4602-0.18258*i |

| | | |
|---|---|---|
| | | $X_4$ = 0.5+0.86603*i<br>$X_5$ = 0.5-0.86603*i |
| $\sqrt[4]{k_2} = \frac{1}{1+\sqrt[8]{k_2}}$ | $x^5 + 2x^3 + x - 1 = 0$ | $X_1$ = 0.56984<br>$X_2$ = -0.5+0.86603*i<br>$X_3$ = -0.5-0.86603*i<br>$X_4$ = 0.21508+1.30714*i<br>$X_5$ = 0.21508-1.30714*i |
| $\frac{1}{\sqrt[4]{k_2}} = 1+\sqrt[8]{k_2}$ | $x^5 - x^4 - 2x^2 - 1 = 0$ | $X_1$ = 1.75488<br>$X_2$ = 0.12256+0.74486*i<br>$X_3$ = 0.12256-0.74486*i<br>$X_4$ = -0.5+0.86603*i<br>$X_5$ = -0.5-0.86603*i |

*See Table S3 for exact root values

**c = 6**

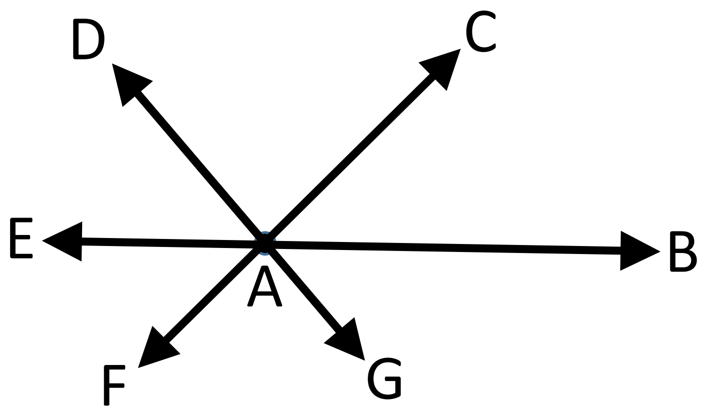


| System Relationships | Correlation with $k_2$ Rate Constant | Value |
|---|---|---|
| $\frac{AG}{AB + AC + AD + AE + AF + AG}$ | $k_2$ | 0.081338 |
| $\frac{AB}{AB+AC+AD+AE+AF+AG} = \frac{AC}{AB+AD+AE+AF+AG} = \frac{AD}{AB+AE+AF+AG} = \frac{AE}{AB+AF+AG} = \frac{AF}{AB+AG} = \frac{AG}{AB}$ | $\sqrt[2]{k_2}$ | 0.285199 |
| $\frac{AB+AC+AD+AE+AF+AG}{AB} = \frac{AB+AD+AE+AF+AG}{AC} = \frac{AB+AE+AF+AG}{AD} = \frac{AB+AF+AG}{AE} = \frac{AB+AG}{AF} = \frac{AB}{AG}$ | $\frac{1}{\sqrt[2]{k_2}}$ | 3.50632 |
| $\frac{AC}{AB} = \frac{AD}{AC} = \frac{AE}{AD} = \frac{AF}{AE} = \frac{AG}{AF}$ | $\sqrt[10]{k_2}$ | 0.77809 |
| $\frac{AB}{AC} = \frac{AC}{AD} = \frac{AD}{AE} = \frac{AE}{AF} = \frac{AF}{AG}$ | $\frac{1}{\sqrt[10]{k_2}} = 1 + \sqrt[2]{k_2}$ | 1.285199 |
| $\frac{AC}{AB+AC+AD+AE+AF+AG} = \frac{AD}{AB+AD+AE+AF+AG} = \frac{AE}{AB+AE+AF+AG} = \frac{AF}{AB+AF+AG} = \frac{AG}{AB+AG}$ | $1 - \sqrt[10]{k_2}$ | 0.22191 |

| | | |
|---|---|---|
| $\frac{AB+AC+AD+AE+AF+AG}{AC} = \frac{AB+AD+AE+AF+AG}{AD} = \frac{AB+AE+AF+AG}{AE} = \frac{AB+AF+AG}{AF} = \frac{AB+AG}{AG}$ | $\frac{1}{1-\sqrt[10]{k_2}} = 1+\frac{1}{\sqrt[2]{k_2}}$ | 4.50633 |
| $\frac{AD}{AB} = \frac{AE}{AC} = \frac{AF}{AD} = \frac{AG}{AE}$ | $\sqrt[5]{k_2}$ | 0.605423 |
| $\frac{AB}{AD} = \frac{AC}{AE} = \frac{AD}{AF} = \frac{AE}{AG}$ | $\frac{1}{\sqrt[5]{k_2}}$ | 1.651737 |

| Polynomial Expressions & Root Values* | | |
|---|---|---|
| $k_2$ | $x^6 - 5x^5 + 10x^4 - 12x^3 - 15x^2 - 11x + 1 = 0$ | **$X_1$ = 0.08134**<br>**$X_2$ = 3.53918**<br>**$X_3$ = -0.50799+0.51585*i**<br>**$X_4$ = -0.50799-0.51585*i**<br>**$X_5$ = 1.19773+2.27876*i**<br>**$X_6$ = 1.19773-2.27876*i** |
| $\sqrt[2]{k_2}$ | $x^6 + 5x^5 + 10x^4 + 10x^3 + 5x^2 + x - 1 = 0$ | **$X_1$ = 0.2852**<br>**$X_2$ = -1.88127**<br>**$X_3$ = -0.32863+0.78485*i**<br>**$X_4$ = -0.32863-0.78485*i**<br>**$X_5$ = -1.37333+0.82964*i**<br>**$X_6$ = -1.37333-0.82964*i** |
| $\frac{1}{\sqrt[2]{k_2}}$ | $x^6 - x^5 - 5x^4 - 10x^3 - 10x^2 - 5x - 1 = 0$ | **$X_1$ = -0.53156**<br>**$X_2$ = 3.50632** |

| | | |
|---|---|---|
| | | $X_3$ = -0.53347+0.32227*i<br>$X_4$ = -0.53347-0.32227*i<br>$X_5$ = -0.45392+1.08406*i<br>$X_6$ = -0.45392-1.08406*i |
| $\sqrt[10]{k_2}$ | $x^6 + x - 1 = 0$ | $X_1$ = 0.77809<br>$X_2$ = -1.13472<br>$X_3$ = 0.62937+0.73576*i<br>$X_4$ = 0.62937-0.73576*i<br>$X_5$ = -0.45106+1.00236*i<br>$X_6$ = -0.45106-1.00236*i |
| $\frac{1}{\sqrt[10]{k_2}} = 1 + \sqrt[2]{k_2}$ | $x^6 - x^5 - 1 = 0$ | $X_1$ = -0.88127<br>$X_2$ = 1.2852<br>$X_3$ = 0.67137+0.78485*i<br>$X_4$ = 0.67137-0.78485*i<br>$X_5$ = -0.37333+0.82964*i<br>$X_6$ = -0.37333-0.82964*i |
| $1 - \sqrt[10]{k_2}$ | $x^6 - 6x^5 + 15x^4 - 20x^3 + 15x^2 - 7x + 1 = 0$ | $X_1$ = 0.22191<br>$X_2$ = 2.13472<br>$X_3$ = 0.37063+0.73576*i<br>$X_4$ = 0.37063-0.73576*i<br>$X_5$ = 1.45106+1.00236*i |

| | | |
|---|---|---|
| | | $X_6$ = 1.45106-1.00236*i |
| $\frac{1}{1-\sqrt[10]{k_2}} = 1+\frac{1}{\sqrt[2]{k_2}}$ | $x^6 - 7x^5 + 15x^4 - 20x^3 + 15x^2 - 6x + 1 = 0$ | $X_1$ = 0.46844<br>$X_2$ = 4.50632<br>$X_3$ = 0.46653+0.32227*i<br>$X_4$ = 0.46653-0.32227*i<br>$X_5$ = 0.54608+1.08406*i<br>$X_6$ = 0.54608-1.08406*i |
| $\sqrt[5]{k_2}$ | $x^6 - 2x^3 - x + 1 = 0$ | $X_1$ = 0.60542<br>$X_2$ = 1.2876<br>$X_3$ = -0.14523+0.92613*i<br>$X_4$ = -0.14523-0.92613*i<br>$X_5$ = -0.80128+0.90424*i<br>$X_6$ = -0.80128-0.90424*i |
| $\frac{1}{\sqrt[5]{k_2}}$ | $x^6 - x^5 - 2x^3 + 1 = 0$ | $X_1$ = 0.77664<br>$X_2$ = 1.65174<br>$X_3$ = -0.54893+0.61947*i<br>$X_4$ = -0.54893-0.61947*i<br>$X_5$ = -0.16526+1.05385*i<br>$X_6$ = -0.16526-1.05385*i |

*There are no exact root values for these polynomials

#### Supplemental Table 3: Exact roots

$$p_1(x) = x^2 - 3x + 1$$

$$x_1 = \frac{1}{2}(3+\sqrt{5}) \approx 2.618033988\,, \qquad x_2 = \frac{1}{2}(3-\sqrt{5}) \approx 0.381966012$$

$$p_2(x) = x^2 + x - 1$$

$$x_1 = \frac{1}{2}(\sqrt{5}-1) \approx 0..6180339880\,, \quad x_2 = -\frac{1}{2}(1+\sqrt{5}) \approx -1.618033988$$

$$p_3(x) = x^2 - x - 1$$

$$x_1 = \frac{1}{2}(\sqrt{5}+1) \approx 1.618033988\,, \quad x_2 = \frac{1}{2}(1-\sqrt{5}) \approx -0.6180339880$$

$$p_4(x) = x^3 - 2x^2 + 5\,x - 1$$

$$\begin{aligned}
x_1 =& -\frac{1}{6}(188+36\sqrt{93})^{1/3} + \frac{22}{3(188+36\sqrt{93})^{1/3}} + \frac{2}{3} \approx 0.2167565713\,,\\
x_2 =& \frac{1}{12}(188+36\sqrt{93})^{1/3} - \frac{11}{3(188+36\sqrt{93})^{1/3}} + \frac{2}{3}\\
&+ \frac{i}{2}\sqrt{3}\left(-\frac{1}{6}(188+36\sqrt{93})^{1/3} - \frac{22}{3(188+36\sqrt{93})^{1/3}}\right) \approx 0.8916217140 - 1.954093394\,i\,,\\
x_3 =& \frac{1}{12}(188+36\sqrt{93})^{1/3} - \frac{11}{3(188+36\sqrt{93})^{1/3}} + \frac{2}{3}\\
&- \frac{i}{2}\sqrt{3}\left(-\frac{1}{6}(188+36\sqrt{93})^{1/3} - \frac{22}{3(188+36\sqrt{93})^{1/3}}\right) \approx 0.8916217140 + 1.954093394\,i
\end{aligned}$$

$$p_5(x) = x^3 + 2\,x^2 + x - 1$$

$$\begin{aligned}
x_1 =& \frac{1}{6}(116+12\sqrt{93})^{1/3} + \frac{2}{3}\left(\frac{1}{(116+12\sqrt{93})^{1/3}} - 1\right) \approx 0.4655712323\\
x_{2,3} =& -\frac{1}{12}(116+12\sqrt{93})^{1/3} - \frac{1}{3}\left(\frac{1}{(116+12\sqrt{93})^{1/3}} + 2\right)\\
&\pm i\,\frac{\sqrt{3}}{2}\left(\frac{1}{6}(116+12\sqrt{93})^{1/3} - \frac{2}{3(116+12\sqrt{93})^{1/3}}\right) \approx -1.232785616 \pm 0.7925519930\,i
\end{aligned}$$

$$p_6(x) = x^3 - x^2 - 2x - 1$$

$$\begin{aligned}
x_1 =& \frac{1}{6}(188 + 12\sqrt{93})^{1/3} + \frac{1}{3}\left(\frac{14}{(188 + 12\sqrt{93})^{1/3}} + 1\right) \approx 2.147899035\,, \\
x_{2,3} =& -\frac{1}{3}\left.\quad \frac{(188 + 12\sqrt{93})^{1/3}}{4} + \frac{7}{(188 + 12\sqrt{93})^{1/3}} - 1\right) \\
& \pm i\,\frac{\sqrt{3}}{6}\left.\quad \frac{(188 + 12\sqrt{93})^{1/3}}{2} - \frac{14}{(188 + 12\sqrt{93})^{1/3}}\right) \approx 0.5739495177 \pm 0.3689894078\,i
\end{aligned}$$

$$p_7(x) = x^3 + x - 1$$

$$\begin{aligned}
x_1 =& \frac{1}{6}(108 + 12\sqrt{93})^{1/3} - \frac{2}{(108 + 12\sqrt{93})^{1/3}} \approx 0.6823278040\,, \\
x_{2,3} =& -\frac{1}{12}(108 + 12\sqrt{93})^{1/3} + \frac{1}{(108 + 12\sqrt{93})^{1/3}} \pm i\,\frac{\sqrt{3}}{2}\left(\frac{(108 + 12\sqrt{93})^{1/3}}{6} + \frac{2}{(108 + 12\sqrt{93})^{1/3}}\right) \\
& \approx -0.3411639019 \pm 1.1615414\,i
\end{aligned}$$

$$p_8(x) = x^3 - x^2 - 1$$

$$\begin{aligned}
x_1 =& \frac{1}{6}(116 + 12\sqrt{93})^{1/3} + \frac{1}{3}\left(\frac{2}{(116 + 12\sqrt{93})^{1/3}} + 1\right) \approx 1.465571232\,, \\
x_{2,3} =& -\frac{1}{12}(116 + 12\sqrt{93})^{1/3} - \frac{1}{3}\left(\frac{1}{(116 + 12\sqrt{93})^{1/3}} - 1\right) \\
& \pm i\,\frac{\sqrt{3}}{6}\left.\quad \frac{(116 + 12\sqrt{93})^{1/3}}{2} - \frac{2}{(116 + 12\sqrt{93})^{1/3}}\right) \approx -0.2327856159 \pm 0.792551993\,i
\end{aligned}$$

$$p_9(x) = x^3 - 3x^2 + 4\,x - 1$$

$$\begin{aligned}
x_1 =& -\frac{1}{6}(108 + 12\sqrt{93})^{1/3} + \frac{2}{(108 + 12\sqrt{93})^{1/3}} + 1 \approx 0.3176721960\,, \\
x_{2,3} =& \frac{1}{12}(108 + 12\sqrt{93})^{1/3} - \frac{1}{(108 + 12\sqrt{93})^{1/3}} + 1 \pm i\,\frac{\sqrt{3}}{2}\left(\frac{(108 + 12\sqrt{93})^{1/3}}{6} + \frac{2}{(108 + 12\sqrt{93})^{1/3}}\right) \\
& \approx 1.341163902 \pm 1.161541400\,i
\end{aligned}$$

$$p_{10}(x) = x^3 - 4x^2 + 3\,x - 1$$

$$x_1 = \frac{1}{6}(188 + 12\sqrt{93})^{1/3} + \frac{1}{3}\left(\frac{14}{(188 + 12\sqrt{93})^{1/3}} + 4\right) \approx 3.147899035\,,$$

$$x_{2,3} = -\frac{1}{12}(188 + 12\sqrt{93})^{1/3} - \frac{1}{3}\left(\frac{7}{(188 + 12\sqrt{93})^{1/3}} - 4\right)$$

$$\pm\, i\,\frac{\sqrt{3}}{6}\quad \frac{(188 + 12\sqrt{93})^{1/3}}{2} - \frac{14}{(188 + 12\sqrt{93})^{1/3}}\Bigg) \approx 0.4260504820 \pm 0.3689894078\,i$$

$$p_{11}(x) = x^5 - 4x^4 + 6\,x^3 + 4\,x^2 + 9\,x - 1$$

$$x_1 = -\frac{1}{3}\quad \frac{(692 + 84\sqrt{69})^{1/3}}{2} - \frac{10}{(692 + 84\sqrt{69})^{1/3}} - 5\Bigg) \approx 0.105441752\,,$$

$$x_{2,3} = \frac{1}{2}(-1 \pm i\,\sqrt{3}) \approx -0.5 \pm 0.8660254040\,i\,,$$

$$x_{4,5} = \frac{1}{3}\left(\frac{(692 + 84\sqrt{69})^{1/3}}{4} - \frac{5}{(692 + 84\sqrt{69})^{1/3}} + 5\right) \pm i\,\frac{\sqrt{3}}{6}\quad \frac{(692 + 84\sqrt{69})^{1/3}}{2} + \frac{10}{(692 + 84\sqrt{69})^{1/3}}\Bigg)$$

$$\approx 2.447279124 \pm 1.869420791\,i$$

$$p_{12}(x) = x^5 + 4\,x^4 + 6\,x^3 + 4\,x^2 + x - 1$$

$$x_1 = \frac{1}{6}(108 + 12\sqrt{69})^{1/3} + \frac{2}{(108 + 12\sqrt{69})^{1/3}} - 1 \approx 0.324717958\,,$$

$$x_{2,3} = \frac{1}{2}(-1 \pm i\,\sqrt{3}) \approx -0.5 \pm 0.8660254040\,i\,,$$

$$x_{4,5} = -\frac{1}{12}(108 + 12\sqrt{69})^{1/3} - \frac{1}{(108 + 12\sqrt{69})^{1/3}} - 1 \pm i\,\frac{\sqrt{3}}{2}\quad \frac{(108 + 12\sqrt{69})^{1/3}}{6} - \frac{2}{(108 + 12\sqrt{69})^{1/3}}\Bigg)$$

$$\approx -1.662358979 \pm 0.5622795125\,i$$

$$p_{13}(x) = x^5 - x^4 - 4x^3 - 6x^2 - 4x - 1$$

$$\begin{aligned}
x_1 =& \frac{1}{6}(388 + 12\sqrt{69})^{1/3} + \frac{26}{3(388 + 12\sqrt{69})^{1/3}} + \frac{2}{3} \approx 3.079595625 \\
x_{2,3} =& \frac{1}{2}(-1 \pm i\sqrt{3}) \approx -0.5 \pm 0.8660254040\, i \\
x_{4,5} =& -\frac{1}{12}(388 + 12\sqrt{69})^{1/3} - \frac{13}{3(388 + 12\sqrt{69})^{1/3}} + \frac{2}{3} \\
& \pm i\,\frac{\sqrt{3}}{2} \left. \quad \frac{(388 + 12\sqrt{69})^{1/3}}{6} - \frac{26}{3(388 + 12\sqrt{69})^{1/3}} \right) \approx -0.5397978113 \pm 0.1825822549\, i
\end{aligned}$$

$$p_{14}(x) = x^5 + x - 1$$

$$\begin{aligned}
x_1 =& \frac{1}{6}(100 + 12\sqrt{69})^{1/3} + \frac{2}{3(100 + 12\sqrt{69})^{1/3}} - \frac{1}{3} \approx 0.7548776667\,, \\
x_{2,3} =& \frac{1}{2}(1 \pm i\sqrt{3}) \approx 0.5 \pm 0.8660254040\, i\,, \\
x_{4,5} =& -\frac{1}{12}(100 + 12\sqrt{69})^{1/3} - \frac{1}{3(100 + 12\sqrt{69})^{1/3}} - \frac{1}{3} \\
& \pm i\,\frac{\sqrt{3}}{6} \left. \quad \frac{(100 + 12\sqrt{69})^{1/3}}{2} - \frac{2}{(100 + 12\sqrt{69})^{1/3}} \right) \approx -0.8774388331 \pm 0.7448617670\, i
\end{aligned}$$

$$p_{15}(x) = x^5 - x^4 - 1$$

$$\begin{aligned}
x_1 =& \frac{1}{6}(108 + 12\sqrt{69})^{1/3} + \frac{2}{(108 + 12\sqrt{69})^{1/3}} \approx 1.324717958\,, \\
x_{2,3} =& \frac{1}{2}(1 \pm i\sqrt{3}) \approx 0.5 \pm 0.8660254040\, i\,, \\
x_{4,5} =& -\frac{1}{12}(108 + 12\sqrt{69})^{1/3} - \frac{1}{(108 + 12\sqrt{69})^{1/3}} \pm i\,\frac{\sqrt{3}}{2} \left. \quad \frac{(108 + 12\sqrt{69})^{1/3}}{6} - \frac{2}{(108 + 12\sqrt{69})^{1/3}} \right) \\
& \approx -0.662358979 \pm 0.5622795125\, i
\end{aligned}$$

$$\boldsymbol{p_{16}(x) = x^5 - 5x^4 + 10\,x^3 - 10x^2 + 6\,x - 1}$$

$$\begin{aligned}
x_1 =& -\frac{1}{6}(100+12\sqrt{69})^{1/3} - \frac{2}{3(100+12\sqrt{69})^{1/3}} + \frac{4}{3} \approx 0.245122333\,,\\
x_{2,3} =& \frac{1}{2}(1 \pm i\sqrt{3}) \approx 0.5 \pm 0.8660254040\,i\,,\\
x_{4,5} =& \frac{1}{12}(100+12\sqrt{69})^{1/3} + \frac{1}{3(100+12\sqrt{69})^{1/3}} + \frac{4}{3}\\
&\pm i\,\frac{\sqrt{3}}{6}\left.\quad \frac{(100+12\sqrt{69})^{1/3}}{2} - \frac{2}{(100+12\sqrt{69})^{1/3}}\right) \approx 1.877438833 \pm 0.7448617670\,i
\end{aligned}$$

$$\boldsymbol{p_{17}(x) = x^5 - 6x^4 + 10\,x^3 - 10x^2 + 5\,x - 1}$$

$$\begin{aligned}
x_1 =& \frac{1}{6}(388+12\sqrt{69})^{1/3} + \frac{26}{3(388+12\sqrt{69})^{1/3}} + \frac{5}{3} \approx 4.079595625\,,\\
x_{2,3} =& \frac{1}{2}(1 \pm i\sqrt{3}) \approx 0.5 \pm 0.8660254040\,i\,,\\
x_{4,5} =& -\frac{1}{12}(388+12\sqrt{69})^{1/3} - \frac{13}{3(388+12\sqrt{69})^{1/3}} + \frac{5}{3}\\
&\pm i\,\frac{\sqrt{3}}{2}\left.\quad \frac{(388+12\sqrt{69})^{1/3}}{6} - \frac{26}{3(388+12\sqrt{69})^{1/3}}\right) \approx 0.460202189 \pm 0.1825822549\,i
\end{aligned}$$

$$\boldsymbol{p_{18}(x) = x^5 + 2\,x^3 + x - 1}$$

$$\begin{aligned}
x_1 =& \frac{1}{6}(44+12\sqrt{69})^{1/3} - \frac{10}{3(44+12\sqrt{69})^{1/3}} + \frac{1}{3} \approx 0.5698402912\,,\\
x_{2,3} =& \frac{1}{2}(-1 \pm i\sqrt{3}) \approx -0.5 \pm 0.8660254040\,i\,,\\
x_{4,5} =& -\frac{1}{12}(44+12\sqrt{69})^{1/3} + \frac{5}{3(44+12\sqrt{69})^{1/3}} + \frac{1}{3}\\
&\pm i\,\frac{\sqrt{3}}{2}\left.\quad \frac{(44+12\sqrt{69})^{1/3}}{6} + \frac{10}{3(44+12\sqrt{69})^{1/3}}\right) \approx 0.2150798545 \pm 1.307141279\,i
\end{aligned}$$

$$p_{19}(x) = x^5 - x^4 - 2x^2 - 1$$

$$\begin{aligned}
x_1 =& \frac{1}{6}(100 + 12\sqrt{69})^{1/3} + \frac{2}{3(100 + 12\sqrt{69})^{1/3}} + \frac{2}{3} \approx 1.754877667\,, \\
x_{2,3} =& \frac{1}{2}(-1 \pm i\,\sqrt{3}) \approx -0.5 \pm 0.8660254040\,i\,, \\
x_{4,5} =& -\frac{1}{12}(100 + 12\sqrt{69})^{1/3} - \frac{1}{3(100 + 12\sqrt{69})^{1/3}} + \frac{2}{3} \\
& \pm i\,\frac{\sqrt{3}}{2} \quad \left. \frac{(100 + 12\sqrt{69})^{1/3}}{6} - \frac{2}{3(100 + 12\sqrt{69})^{1/3}} \right) \approx 0.1225611669 \pm 0.7448617670\,i
\end{aligned}$$
